# The brain integrates proprioceptive information to ensure robust locomotion

**DOI:** 10.1101/2021.04.28.441796

**Authors:** Alessandro Santuz, Olivier D. Laflamme, Turgay Akay

## Abstract

Robust locomotion relies on information from proprioceptors: sensory organs that communicate the position of body parts to the spinal cord and brain. Proprioceptive circuits in the spinal cord are known to regulate locomotion in challenging environments. Yet, the regulatory importance of the brain remains less clear. Here, through mouse genetic studies and in vivo electrophysiology, we examined the role of the brain in integrating proprioceptive information during locomotion. The systemic removal of proprioceptors left the animals in a constantly perturbed state, similar to that observed during mechanically perturbed locomotion in wild type and characterised by longer and less accurate activation patterns. In contrast, after surgically interrupting the ascending proprioceptive projection to the brain through the dorsal column pathway, wild-type mice showed normal walking behaviour, but lost the ability to respond to external perturbations. Our findings provide direct evidence of a pivotal role for ascending proprioceptive information in achieving safe locomotion.

## Introduction

Animals constantly move in complex environments. Locomotor actions often dictate the safe execution of challenging activities such as chasing prey, escaping danger, exploring new territories. In vertebrates, the generation of rhythmic and patterned activities like those needed for locomotion is in part achieved through neuronal networks located in the spinal cord: the central pattern generators (Brown, 1911). The higher order integration of spatiotemporal information happening in the cerebral cortex and subcortical regions is known to be important for initiating and halting locomotion or regulating its speed (Cignetti et al., 2017; Karadimas et al., 2019; Roseberry and Kreitzer, 2017; Rossignol et al., 2006). Yet, when somatosensory feedback is removed and supraspinal pathways are left intact, the motor commands generated by the central pattern generators alone are not sufficiently robust to allow for safe locomotion in the presence of external perturbations (Akay, 2020; Grillner and El Manira, 2020; Santuz et al., 2019).

In mammals, information about body position in space and relative to the body itself (i.e., “proprioception”) is conveyed by mechanosensory neurons found within muscles, tendons and joints (Dietz, 2002). These neurons bear information from the muscle spindles, Golgi tendon organs (GTOs) and joint receptors. While not much is known about the function of the latter (Tuthill and Azim, 2018), feedback from muscle spindles (group Ia and II afferent fibres) and GTOs (group Ib) undoubtedly plays a role in mammalian locomotion (Grillner and El Manira, 2020; Pearson, 1995). In cats, mice, humans and other mammals, these proprioceptive afferents are pivotal for setting the timing of the gait cycle and facilitating the switch between different phases, such as propulsion and swing (Rossignol et al., 2006). In mice, the selective and systemic removal of proprioceptive afferents can disrupt the locomotor patterns (Akay et al., 2014; Takeoka and Arber, 2019) and the regulation of muscle activity needed to locomote at different speeds (Mayer et al., 2018) or to cope with perturbations (Santuz et al., 2019). Part of the proprioceptive information is elaborated locally in the spinal cord (spinal sensory processing), but there are three, topographically well distinct, main pathways that convey proprioceptive feedback from the hindlimb to brain structures (supraspinal sensory processing): the dorsal and ventral spinocerebellar tracts (Bosco and Poppele, 2001) and the dorsal column-medial lemniscus (DCML) pathway (Niu et al., 2013). The dorsal spinocerebellar tract originates from the Clarke’s column that receives somatosensory information through the dorsal column and conveys it to the cerebellum through the lateral funiculi; it is thought to be the major carrier of proprioceptive and exteroceptive (i.e., touch and pressure) signals directly to the cerebellum from the hindlimb (Hantman and Jessell, 2010; Lundberg, 1964). Similarly, the ventral spinocerebellar tract relays some proprioceptive information to the cerebellum via the lateral funiculi, but ventral relatively to the dorsal spinocerebellar pathway (Chalif et al., 2022; Jankowska et al., 2010). Contrary to the spinocerebellar tracts, the DCML pathway transmits to the somatosensory cortex via the brainstem and it mostly conveys signals from proprioceptors and touch exteroceptors (Niu et al., 2013). While the neural circuitry underlying sensory feedback from muscle spindles and GTOs in the dorsal column is well investigated, the importance of higher centres in integrating proprioceptive information during locomotion is still obscure (Akay, 2020).

Here, by using a combination of mouse genetics, in vivo electrophysiology, spinal lesion models and computational neuroscience (Fig. 1), we set out to explore how relevant is the supraspinal integration of proprioceptive feedback via DCML pathway during murine locomotion. We posited that not only spinal but also supraspinal sensory processing must be involved in the tuning of motor control when locomotion becomes challenging. During walking, we administered random mediolateral and anteroposterior perturbations to four groups of animals by means of sudden lateral displacements or accelerations of the treadmill’s belt. A first group of wild-type served as control. Then, we added to the setup a genetically modified strain (*Egr3^-/-^*) in which muscle spindles regress immediately after birth. To control for potential adaptation to the lack of sensory feedback during development, we included a third group in which all proprioceptors could be ablated systemically and acutely in adult age (*PV^Cre^::Avil^iDTR^*). Lastly, we surgically lesioned the dorsal column to disrupt proprioceptive information reaching the brain through the DCML and partly through the dorsal spinocerebellar pathways in wild type animals to isolate part of the proprioceptive information travelling from the hindlimb to the brain, leaving the local spinal circuitry intact. By means of high-speed video recordings, electromyography (EMG) and an analysis framework based on linear and nonlinear tools, we show that supraspinal processing of proprioceptive feedback becomes crucial if locomotion happens is challenged by external perturbations.

**Fig. 1.**
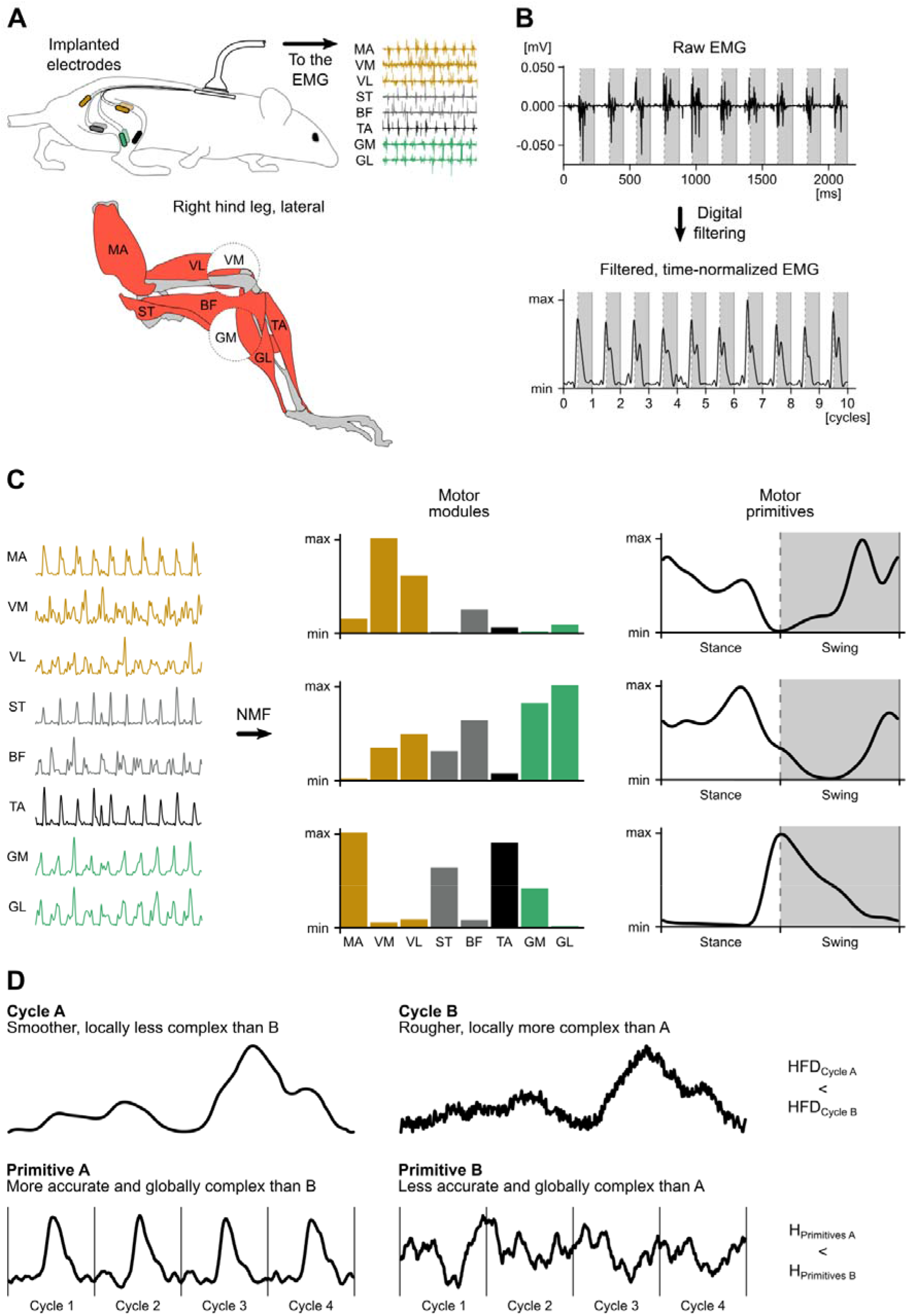
Schematic representation of the muscle synergies approach and following fractal analysis. (**A**) Eight muscles were implanted in the right hindlimb. (**B**) Raw EMG was filtered and time- and amplitude-normalized. (**C**) Filtered and normalized EMG was factorized with non-negative matrix factorization and classified into motor modules (time-invariant muscle weightings) and motor primitives (time-dependent synergistic coefficients). (**D**) The Higuchi’s fractal dimension (HFD, top row) is a measure of local complexity and it increases together with the “roughness” of motor primitives at a single cycle level (thus the term “local”). The Hurst exponent (H, bottom row) is a measure of global complexity and it increases if the “accuracy” of motor primitives decreases across several gait cycles (thus the term “global”).

## Results

### Wild-type mice show robust locomotion despite perturbations

First, we investigated the effects of external perturbations on locomotion in wild-type mice (Fig. 2A, Fig. 3, movie S1). The perturbation protocol (Fig. S1) increased the cadence, or number of steps per minute (Fig. S3) and the step-to-step variability of the hip joint angle at touchdown, as shown by the Poincaré maps (Fig. 3A, Fig. S4). Additionally, perturbations reduced the amount of flexion of the knee- and ankle-joints in the early stages of the swing phase (Fig. 3B). The root mean square values of the EMG (RMS_EMG_), indicator of the signal’s amplitude, was on average slightly higher during perturbed locomotion, but only when perturbations were only mediolateral and not random (i.e., every 2 s; Fig. 3C, Fig. S5). When perturbations were randomly administered, the effect on the RMS_EMG_ was negligible. To uncover the modular structure of muscle activations, we decomposed EMG activity into muscle synergies via non-negative matrix factorization (Bizzi et al., 2008; Lee and Seung, 1999). Muscle synergies are represented by a set of time-invariant muscle weightings (or motor modules, describing the relative contribution of each muscle to a specific synergy) and a set of time-dependent activation patterns (or motor primitives). Three synergies were sufficient to reconstruct the original EMG signals of both unperturbed and perturbed walking (Fig. 3D). The obtained motor modules and motor primitives described three main phases of the gait cycle: 1) the weight acceptance, with the major contribution of knee extensors; 2) the propulsion, mostly involving the ankle extensors; 3) and the swing, characterized by the contribution of hip abductors and knee and ankle flexors.

**Fig. 2.**
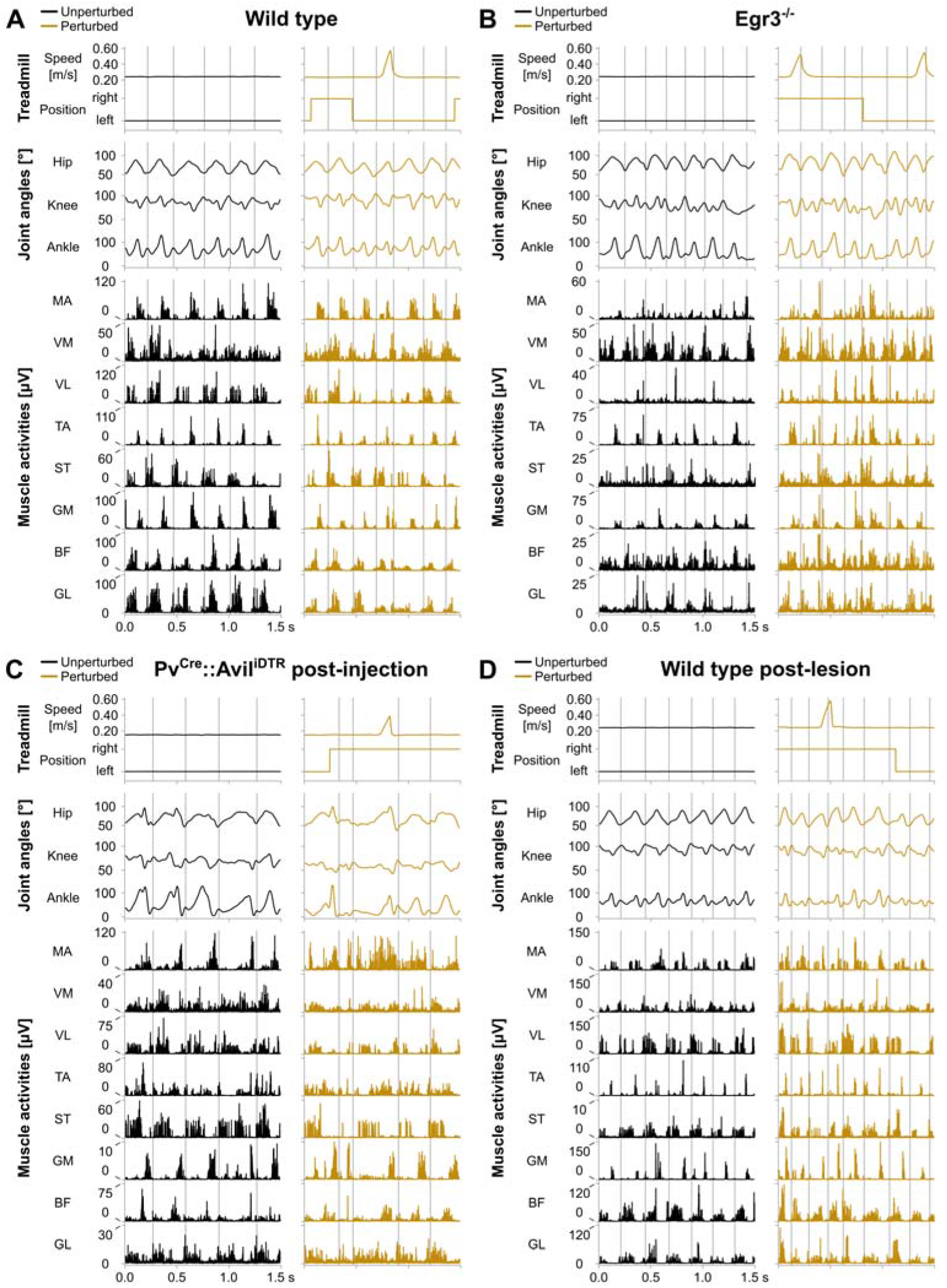
Hindlimb joint angles and muscle activities during unperturbed and perturbed locomotion. (**A**) Treadmill speed and mediolateral position, joint angles and rectified raw muscle activities of a representative wild-type mouse, without time-normalisation of the gait cycles. Vertical lines represent the touchdown of each gait cycle. Muscle abbreviations are reported in the caption of Fig. 3. (**B**) The same as in (**A**), but for a representative *Egr3*^-/-^ mouse. (**C**) The same as in (**A**), but for a representative *Pv^Cre^::Avil^iDTR^* mouse after DTX injection. (**D**) The same **as** in (**A**), but for a representative wild-type mouse after surgical lesion of the DCML pathway in the spinal cord.

**Fig. 3.**
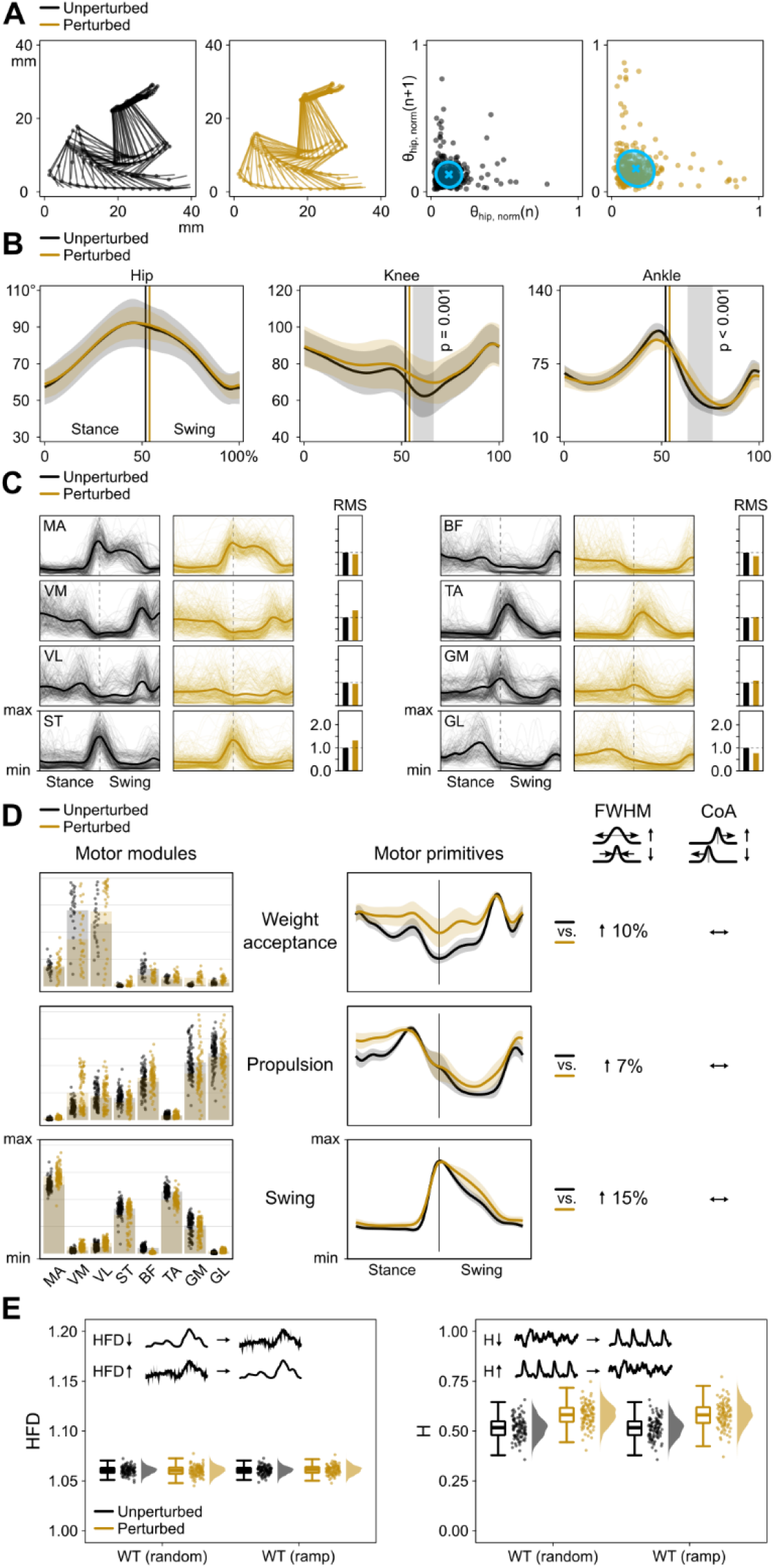
Gait performance parameters of wild-type mice during unperturbed and perturbed locomotion. (**A**) Stick diagrams of one representative animal (left) and Poincaré maps (right) of the hip angle θ_hip_ at touchdown (n) and (n+1) of all animals, with descriptor ellipse (see methods). (**B**) Average joint angles (thick lines) and standard deviation (shaded bands) for all animals. Vertical shaded areas denote differences detected by statistical parametric mapping. (**C**) Average electromyographic activity (thick lines) of the eight recorded muscles and individual trials (thin lines) from all five mice. RMS is normalized to unperturbed walking. Muscle abbreviations: MA = *gluteus maximus*, VM = *vastus medialis*, VL = *v. lateralis*, ST = *semitendinosus*, BF = *biceps femoris*, TA = *tibialis anterior*, GM = *gastrocnemius medialis*, and GL = *g. lateralis*. (**D**) Motor modules and primitives of the three bootstrapped muscle synergies. Modules are presented on a normalized y-axis base. For simplifying the graphs, each point represents 10 nearest neighbours of the 1000 bootstrapped trials. For the motor primitives, the x-axis full scale represents the averaged step cycle (stance and swing normalized to the same amount of points) and the y-axis the normalized amplitude. Full width at half maximum (FWHM) and centre of activity (CoA) of motor primitives are reported schematically (right). (**E**) Boxplots describing the Higuchi’s fractal dimension (local complexity HFD or “roughness”) and Hurst exponent (global complexity H or “accuracy”) of the bootstrapped primitives for two perturbation protocols: random (Fig. S1) and one mediolateral displacement every 2 s (ramp). Raw data points (each point represents 10 nearest neighbours of the 1000 bootstrapped trials) and their density estimates are presented to the right side of each boxplot.

During perturbed locomotion, compared to unperturbed walking, the timing of motor primitives did not undergo any noteworthy shift, as measured by the centre of activity (Fig. 3D, Fig. S8). However, all primitives were longer active, as shown by the increased full width at half maximum (Fig. 3D, Fig. S7). To further investigate the reasons for this widening, we expanded the analysis towards nonlinear metrics that could give more information about, for instance, an increased variability or regularity of the primitives that might have caused the longer activation profile of the primitives. The global (Hurst exponent) but not the local (Higuchi’s fractal dimension) complexity of motor primitives was affected by perturbations in wild-type mice (Fig. 3E, Fig. S9). The local complexity can be seen as a measure of “roughness” (or noise content) in the signal within each gait cycle, while the global complexity is a measure of how accurate each cycle’s activation motifs are, when compared to the others in the same trial (Fig. 1D). Motor primitives were less accurate and complex (i.e., higher Hurst exponent) when the animals were challenged by perturbations. An outcome that might explain the widening found with the analysis of the full width at half maximum. These first observations in wild-type animals could show that perturbations to locomotion in intact animals elicited: a) higher variability and less flexion of joint angles; b) similar amplitude of EMG signals; and c) wider, less accurate and globally less complex motor primitives.

### Feedback from muscle spindles regulates locomotion

Then, we set out to study the aftermath of genetic removal of one class of proprioceptors: the muscle spindles. We used *Egr3*^-/-^ mice (Tourtellotte and Milbrandt, 1998), a model in which muscle spindles regress after birth (Fig. 2B, movie S2). In those mutant animals, the number of steps per minute was higher than in wild type and the swing phase was shorter (Fig. S3). Moreover, in this group the kinematics were generally not affected by the presence of external perturbations, but the variability of hip joint angle at touchdown was higher than in wild type (Fig. 4A and B, Fig. S4). Contrary to wild-type animals, *Egr3*^-/-^ did not increase the RMS_EMG_ when exposed to perturbations (Fig. 4C, Fig. S5). Yet, they showed increased duration of the weight acceptance and swing primitives when compared to their wild-type littermates and the main activity shifted later in time in the swing primitive in the absence of perturbations (Fig. 4D, Fig. S6, Fig. S7, Fig. S8). When exposed to perturbations though, mutants only produced a shorter and earlier activation motif in the muscle synergy for the weight acceptance (Fig. 4D, Fig. S6, Fig. S7, Fig. S8). As in wild-type animals, the roughness (i.e., the local complexity) of motor primitives was not affected by perturbations and was not different from that of controls (Fig. 4E, Fig. S9). However, the activation patterns in both normal and perturbed locomotion were as accurate (i.e., same global complexity) as those of perturbed locomotion in wild type (Fig. 4E, Fig. S9). In short, this second part of the experiment revealed that the systemic lack of muscle spindles produced: a) increased variability of kinematics; b) the inability to modulate the hindlimb kinematics or EMG amplitude in the presence of perturbations; and c) wider, less accurate (i.e., globally less complex) motor primitives when compared to wild type, with no nuances when comparing unperturbed and perturbed walking.

**Fig. 4.**
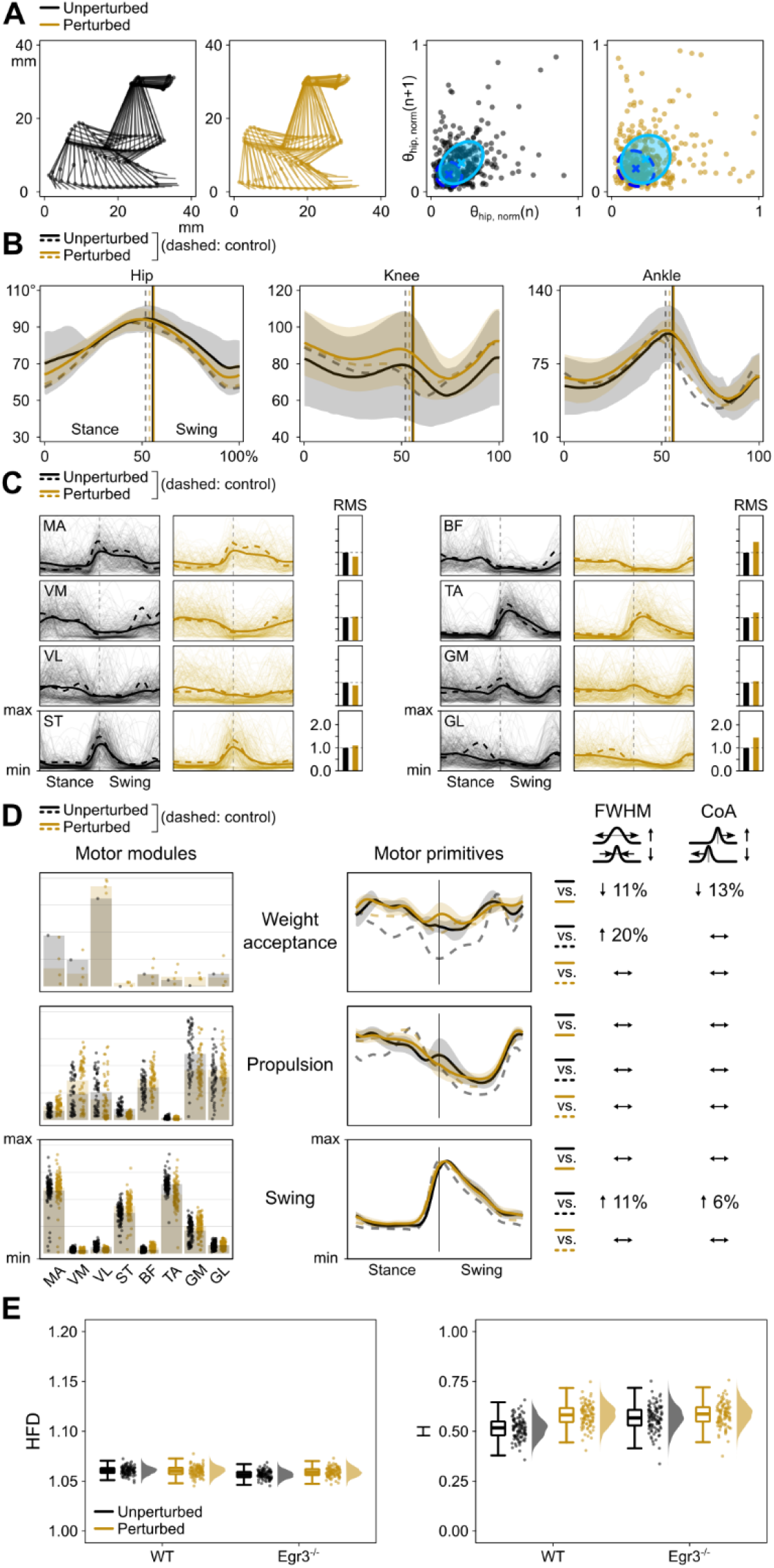
Gait performance parameters of *Egr3*^-/-^ mice during unperturbed and perturbed locomotion. (**A**) The descriptor ellipses in the Poincaré maps are overlaid to those of wild type (Fig. 3). (**B** to **D**) Control is wild type. The RMS in (**B**) is normalized to unperturbed walking of *Egr3*^-/-^. See caption of Fig. 3 for all other details.

### Acute ablation of proprioceptors severely disrupts locomotion

The genetic removal of muscle spindles from birth can however raise the question of whether the animals could adapt to the partial lack of sensory feedback during the first weeks of life. Moreover, the force-sensitive GTOs are known to be important players in the control of locomotion (Donelan et al., 2009), but the *Egr3*^-/-^ model only targets muscle spindles. To tackle these two potential issues, we acutely ablated both muscle spindles and GTOs in adult mice by systemic injection of diphtheria toxin in *PV^Cre^::Avil^iDTR^* mice, as previously done by Takeoka and Arber (Takeoka and Arber, 2019). Here, by immunohistochemistry after each experiment, we confirmed their observation that diphtheria toxin injection is efficient in removing proprioceptive afferent neurons in this mouse line (Fig. 5).

**Fig. 5.**
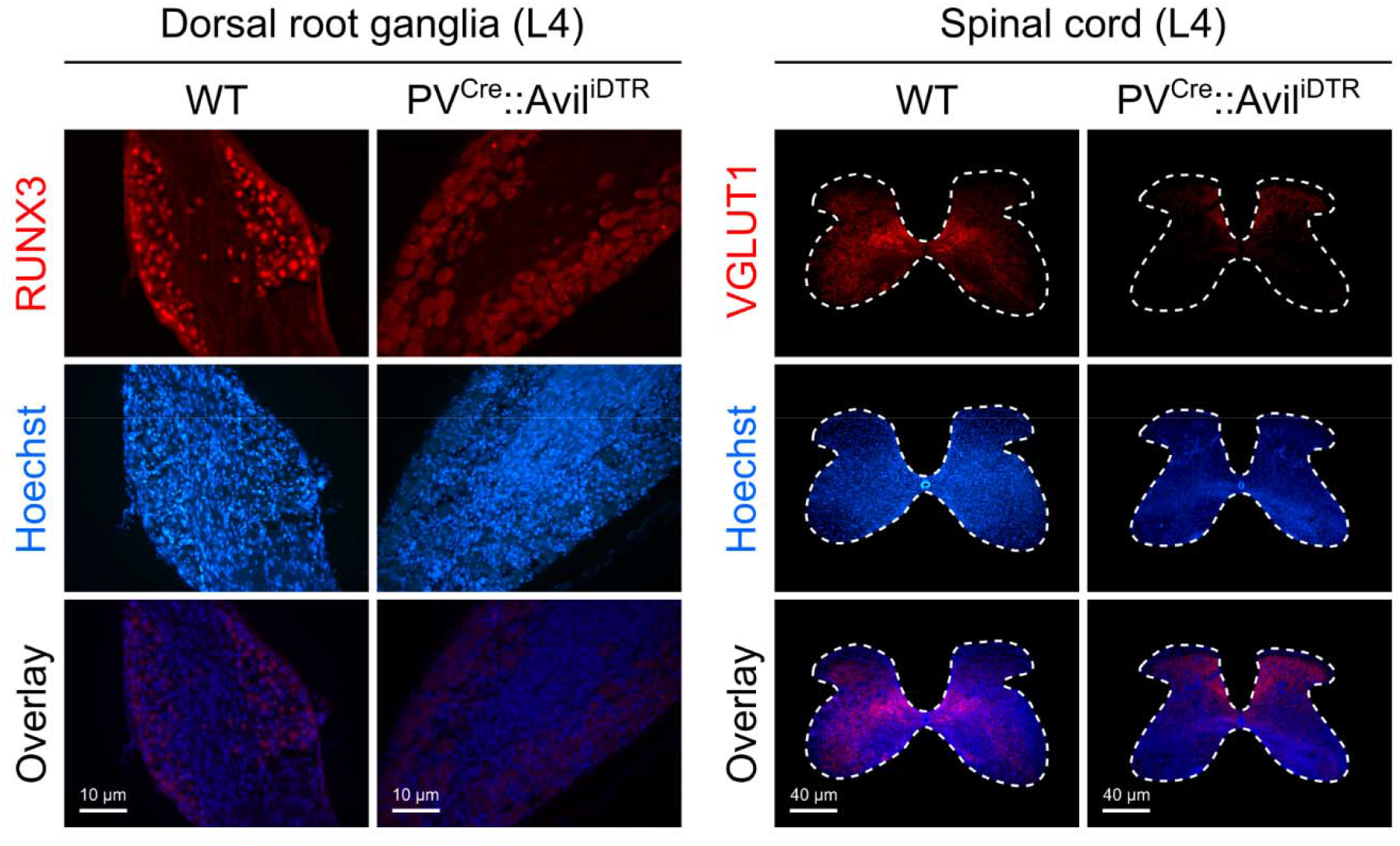
Acute ablation of proprioceptive afferents with diphtheria toxin (DTX). Dorsal root ganglia (left) and spinal cord (right) sections taken at the fourth lumbar spinal segment (L4) in wild-type (WT) and mutant (*PV^Cre^::Avil^iDTR^*) mice. Runt-related transcription factor 3 (RUNX3) is selectively expressed in intact proprioceptive sensory neurons located in dorsal root ganglia. Vesicular glutamate transporter 1 (VGLUT1) is expressed in central sensory terminals. After DTX injection in *PV^Cre^::Avil^iDTR^* mice, the proprioceptive neuron terminals are eliminated at all spinal levels, leaving the fewer in number corticospinal and non-proprioceptive sensory terminals unaffected.

Before injection, these mice behaved largely similar to wild-type animals (Fig. S10). After injection however, they underwent major degradation of sensory neurons resulting in severely disrupted kinematics, largely independently on the presence of perturbations (Fig. 6A and B, Fig. 2C, movie S3). Specifically, the step-to-step variability of the hip joint angle at touchdown increased dramatically as compared to pre-injection recordings (Fig. S4) and the ankle joint was less flexed throughout almost all of the stance phase (Fig. 6B). Moreover, there was a large effect of sensory ablation on the stance and swing time variability (Fig. S2) and duration (Fig. S3). The RMS_EMG_ was on average higher and also unaffected by perturbations after injection (Fig. 6C, Fig. S5). While the number and function of muscle synergies (Fig. 6D) were not influenced by acute sensory ablation, the timing of motor primitives was indeed. When comparing pre- and post-injection recordings, we detected a main effect on the timing of main activation, which was shifted to the right and of shorter duration in the propulsion synergy (Fig. 6D, Fig. S6, Fig. S7, Fig. S8). All the remaining main effects were small and interactions had large effect size in all synergies for both duration and main peak activity, a sign of cross-over interaction or opposite effects in the two groups (Fig. S7, Fig. S8). Similar to what we observed in *Egr3*^-/-^ mice, the now acute disruption of feedback from proprioceptors produced less accurate and complex activation patterns (i.e., higher Hurst exponent), once more making unperturbed and perturbed locomotion numerically indistinguishable (Fig. 6E, Fig. S9). In addition, we found that the sensory ablation was followed by a large increase in the roughness of motor primitives (Fig. 6E, Fig. S9). In summary, the acute ablation of both muscle spindles and GTOs induced: a) increased kinematic variability and EMG amplitude and the inability to modulate them in the presence of perturbations; and b) rougher, less accurate and globally less complex motor primitives, with substantial reorganization of timing.

**Fig. 6.**
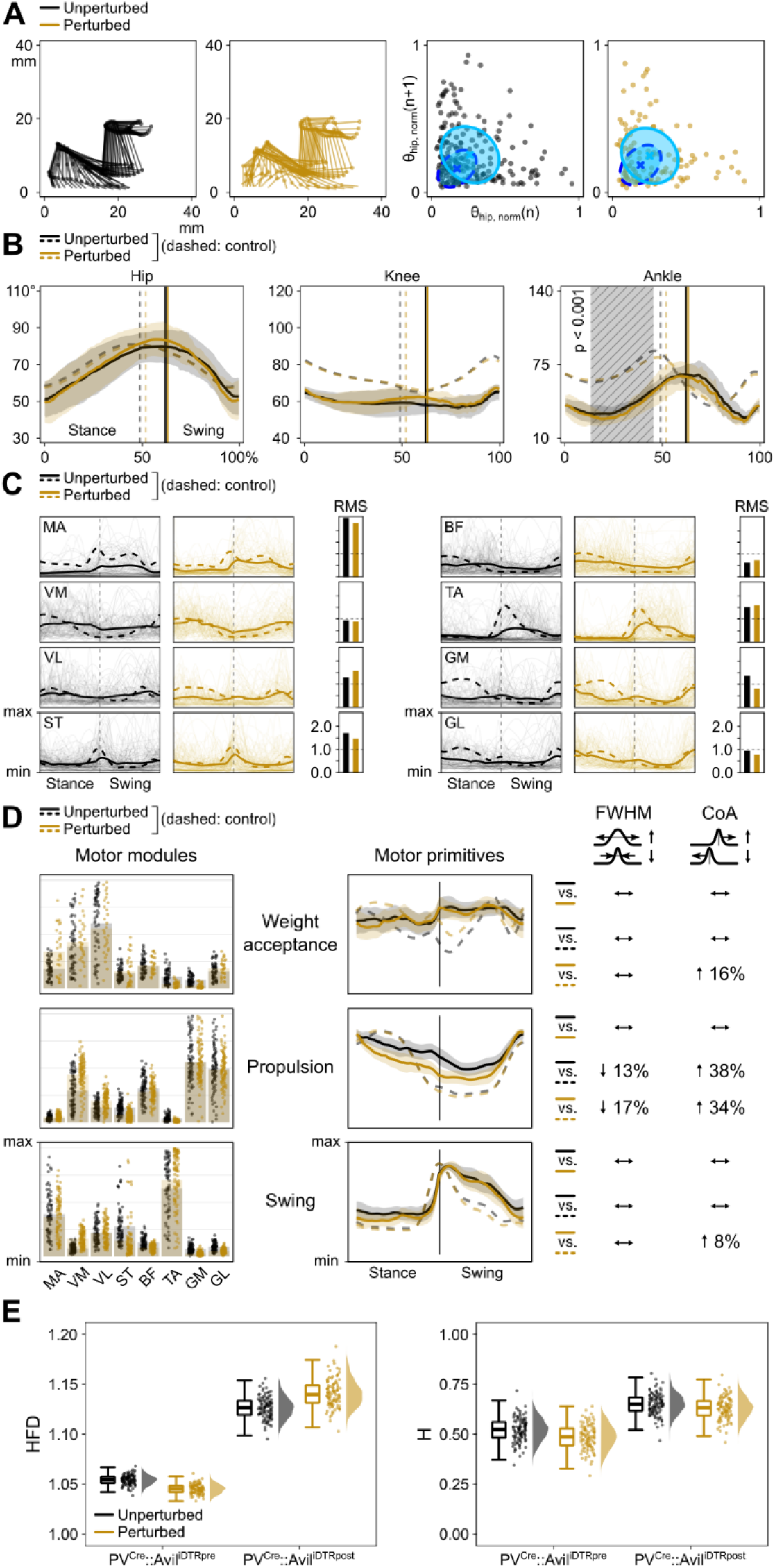
Gait performance parameters of *PV^Cre^::Avil^iDTR^* mice during unperturbed and perturbed locomotion. (**A**) The descriptor ellipses in the Poincaré maps are overlaid to those of pre-DTX injection recordings. (**B** to **D**) Control is pre-injection. The RMS in (**B**) is normalized to unperturbed walking recorded pre-injection. See caption of Fig. 3 for all other details.

### Pivotal role of the brain in controlling challenging locomotion

The neurophysiological changes genetically induced in *Egr3*^-/-^ and *PV^Cre^::Avil^iDTR^* animals were affecting both the spinal as well as the ascending proprioceptive information conduction, thus hindering any reasoning on the specific contribution of supraspinal proprioceptive integration to locomotion. To overcome this last hurdle in our experimental design, we surgically lesioned the dorsal column in wild-type mice to disrupt proprioceptive information projecting to the brain through the DCML and partly the dorsal spinocerebellar pathways (Fig. 7). After lesion, the animals did not show notable changes in either the kinematics or EMG amplitude when compared to pre-lesion recordings (Fig. 8A, B and C, Fig. 2D, Fig. S2, Fig. S3, Fig. S4, Fig. S5, movie S4). The number and function of synergies was not affected by the lesion either (Fig. 8D). Yet, the timing of motor primitives underwent interesting modifications. First, in lesioned animals there was a negligible effect of perturbations on both the duration (Fig. S7) and the main activity timing (Fig. S8), contrarily to what happened before the lesion. Moreover, the motor primitives during both unperturbed and perturbed locomotion were similarly accurate as those of unperturbed walking in intact animals (Fig. 8E, Fig. S9). Surprisingly, three out of five animals would not manage sudden accelerations of the treadmill’s belt, interrupting locomotion immediately after the stimulus was administered. This outcome did not correlate with the extent of the lesion. In summary, with this last part we could show that inhibiting the flow of proprioceptive information to the brain in wild-type animals caused minimal changes to unperturbed locomotion but the loss of ability to interact with perturbations, often resulting in the abrupt termination of locomotion.

**Fig. 7.**
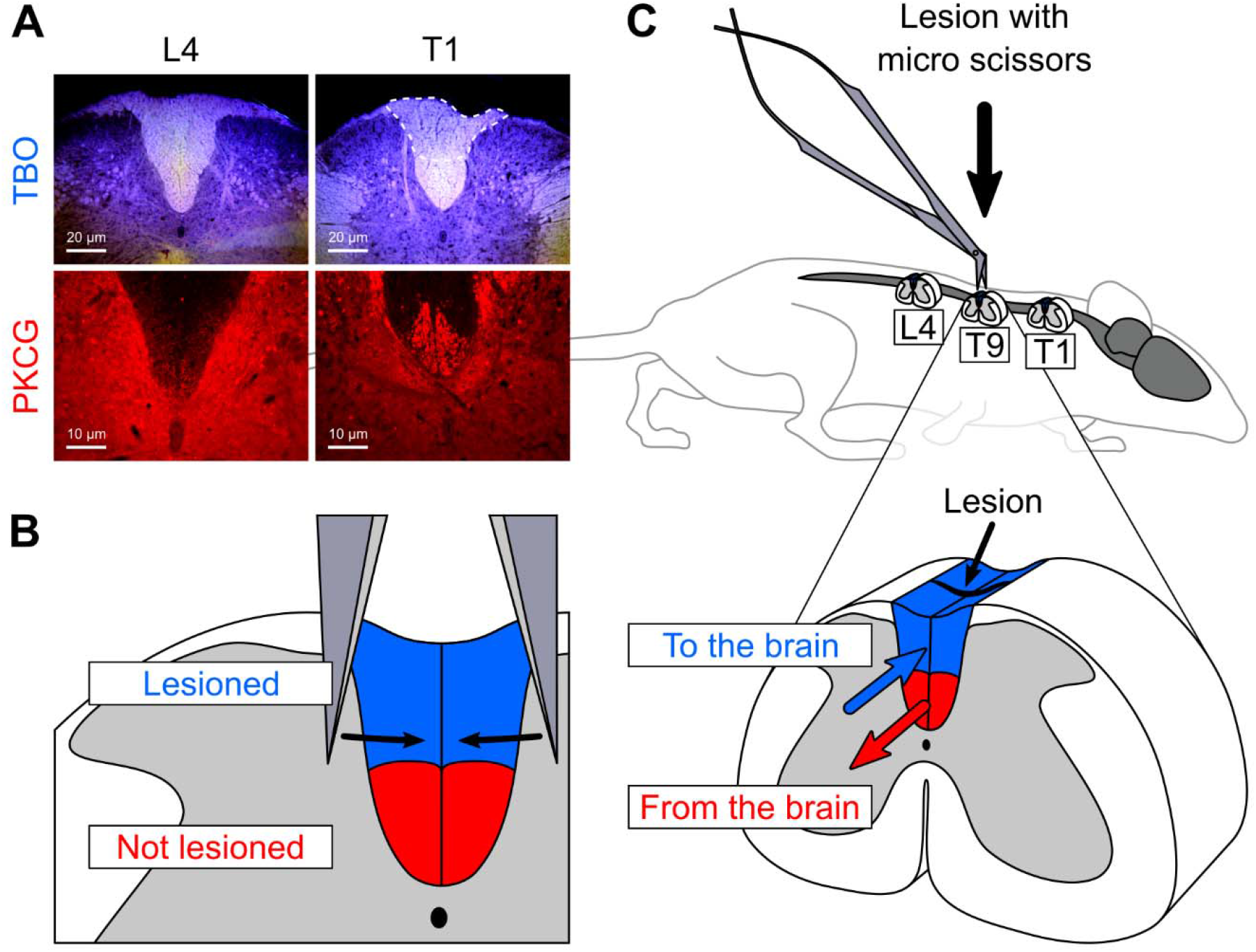
Lesion of the dorsal column-medial lemniscus (DCML) pathway leaves corticospinal tract intact. (**A**) After toluidine blue (TBO) staining, lesioned afferent axons resulted in damaged tissue rostral (first thoracic spinal segment T1) but not caudal (fourth lumbar spinal segment L4) with respect to the lesion. Protein kinase C gamma (PKCG) is expressed in the axons of the corticospinal tract (CST) and can be used to evaluate the integrity of the latter after surgery. (**B**) The surgery was aimed at the lesion of the DCML (most dorsal portion of the dorsal column), leaving the CST (most ventral portion of the dorsal column) intact. (**C**) The lesion was carried out at the ninth thoracic spinal segment (T9). Sections at L4 and T1 were used to evaluate the success of the surgery.

**Fig. 8.**
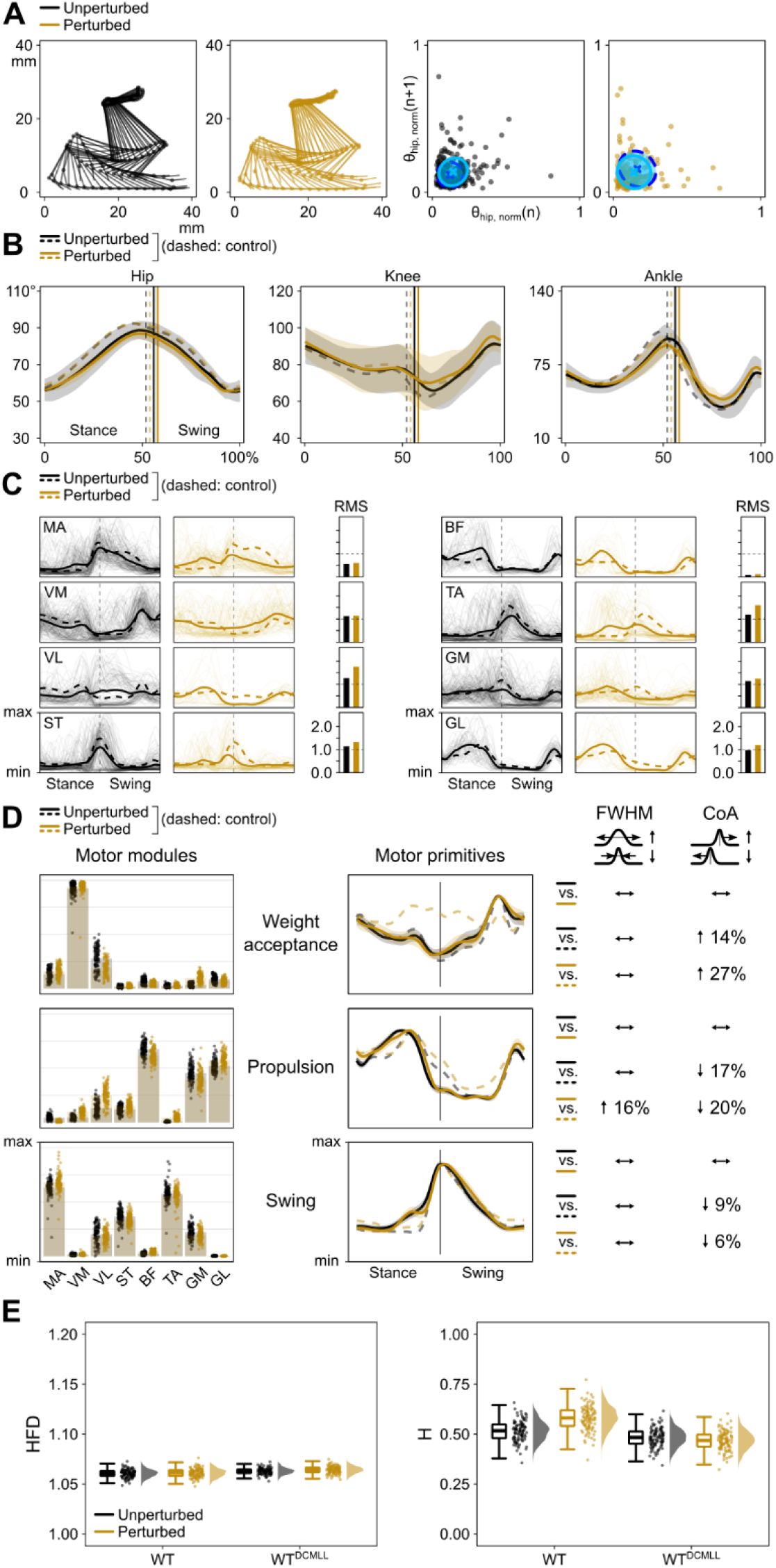
Gait performance parameters of wild-type mice during unperturbed and perturbed locomotion after DCML pathway lesion. (**A**) The descriptor ellipses in the Poincaré maps are overlaid to those of wild type, prelesion (Fig. 3). (**B** to **D**) Control is pre-lesion. The RMS in (**B**) is normalized to unperturbed walking recorded prelesion (Fig. 3). See caption of Fig. 3 for all other details.

## Discussion

We designed a simple experiment to assess whether the murine brain integrates proprioceptive information form the hindlimb through the DCML pathway during locomotion. By using a mix of mouse genetics, in vivo electrophysiology and a spinal lesion model, we showed that supraspinal integration of signals form proprioceptors is crucial for locomotion in the presence of perturbations. This implies that spinal proprioceptive circuits alone are not sufficient to guarantee effective responses to external perturbations during locomotion.

### The systemic removal of proprioceptors undermines the robustness of locomotor control

As previously reported (Mayer and Akay, 2018; Santuz et al., 2019), wild-type mice are exceptional at coping with external perturbations during locomotion. Our results showed that the neural strategies adopted to robustly overcome sudden mediolateral displacements and accelerations produced lower joint flexion and increased kinematic variability. Despite similar amplitudes of the EMG signals, the synergistic activation patterns (i.e., muscle synergies) were tuned after perturbations. In particular, as usually found in both mice (Santuz et al., 2019) and humans (Cappellini et al., 2016; Martino et al., 2014; Santuz et al., 2018), time-dependent activation motifs (i.e., motor primitives) were wider relative to the gait cycle in the presence of perturbations, reflecting an increase in the robustness of motor control. In addition, motor primitives were substantially less accurate and complex in perturbed than in normal walking, similar to what we previously found in mice (Santuz and Akay, 2020) and humans (Santuz et al., 2020a) locomoting in a similar experimental environment. As expected, most of these observations changed when feedback from proprioceptors was either genetically inhibited from birth or acutely removed in adult mice.

The major disruption of locomotor patterns in *Egr3*^-/-^ (Tourtellotte and Milbrandt, 1998) and *PV^Cre^::Avil^iDTR^* (Takeoka and Arber, 2019) mice is well known. *Egr3*^-/-^ lack muscle spindles from birth and typically show gait ataxia (Tourtellotte and Milbrandt, 1998), especially to be seen in the timing of the ankle joint flexors during the swing phase (Akay et al., 2014). In contrast, *PV^Cre^::Avil^iDTR^* mice are born intact and only after acute ablation of feedback from both muscle spindles and GTOs they undergo a swift and irreversible degradation of locomotor capabilities (Takeoka and Arber, 2019) that is by far worse than that observed in *Egr3*^-/-^ mutants. Here, we built up on previous research further characterising kinematics and muscle activation patterns during locomotion with and without perturbations. We found that both mouse lines showed kinematics and muscle activity patterns that were numerically undistinguishable when comparing unperturbed and perturbed locomotion, showing the lack of ability to adapt to external perturbations. Moreover, the vast majority of analysed metrics had values that resembled those found in the wild type during perturbed locomotion. When compared to controls (i.e., wild type for *Egr3*^-/-^ and pre-ablation recordings in *PV^Cre^::Avil^iDTR^*), the outcomes were all similar: increased variability of kinematics and wider, less accurate motor primitives, independently on whether perturbations were administered or not. This indicates an inherently disrupted control of locomotion due to the systemic lack of feedback from proprioceptors, independently on the external challenges.

While *Egr3*^-/-^ and *PV^Cre^::Avil^iDTR^* mice shared many of the outcomes, a few peculiarities were found in the latter that were not visible in the former. The dramatic effects on the timing of propulsion in *PV^Cre^::Avil^iDTR^*, which were not to be seen in *Egr3*^-/-^ mice, might be explained by the absence of feedback from GTOs. Typically, GTOs outnumber muscle spindles by 80% (Prochazka, 2015). Moreover, experiments in the cat during unperturbed walking (Donelan et al., 2009) showed that force feedback is responsible for around one third of the total muscle activity in the flexors of the ankle, muscles that are mostly contributing to the propulsion synergy. This share increases with the task demand (Donelan et al., 2009) and/or the length at which muscle fascicles work (Donelan and Pearson, 2004) and could thus explain the effect of spindle and GTO removal on the timing of the propulsion synergy. Moreover, the sensory ablation in *PV^Cre^::Avil^iDTR^* mice was followed by a large increase in the roughness of motor primitives. However, this might be due to the known effect of speed on the Higuchi’s fractal dimension (Santuz et al., 2020b): after injection, the animals could not cope with pre-injection speeds and had to be tested at half those speeds (0.15 instead of 0.25 and 0.30 m/s), also resulting in longer cycle times and lower cadence (Fig. S2, Fig. S3).

### Ascending proprioceptive pathways in the dorsal column carry crucial information for locomotion in challenging conditions

Surgical lesion of dorsal column in wild-type mice allowed us to interrupt a major part of the proprioceptive information flowing from the hindlimb to the brain, leaving all spinal circuits intact. The murine DCML pathway is composed, caudal to the sixth thoracic segment, by two tracts in the dorsal column of the white matter (Niu et al., 2013): the medial gracile fasciculus (dorsal region) and the corticospinal tract (ventral region). Given its extremely dorsal location (Watson and Harrison, 2012), the dorsal column could be easily exposed during surgery. The lesion would disrupt the ascending information to the brainstem from proprioceptors and exteroceptive touch sensors, but mostly from hind-limb proprioceptors when surgery is carried out in the lower thoracic spinal segments (Niu et al., 2013). This surgery, however, would partially or fully damage the corticospinal tract (mainly connecting the motor cortex to the spinal cord) as its projection is just ventral to the ascending projections in the dorsal column (Lieu et al., 2013). Moreover, the spinocerebellar tract would be left intact since this pathway projects through the lateral funiculus to the cerebellum (English, 1985; Mann, 1973).

In unperturbed locomotion, the lesion did not produce notable changes in the kinematics, EMG amplitude or motor primitive timings compared to data collected before surgery in the same animals. This is in agreement with previous findings after dorsal column lesion in the cat (English, 1985, 1980). Yet, a striking fact emerged when administering external perturbations: the animals behaved almost identically as in the unperturbed state, leading to similarities in the vast majority of the analysed outcomes. In other words, lesioned animals were unable to discern the perturbed state from the unperturbed and this can be confirmed by recalling the observations done in proprioception-deficient mutants. Both *Egr3*^-/-^ and *PV^Cre^::Avil^iDTR^* mice showed some adaptation to the partial or total lack of proprioceptive information, which was visible in the constant, perturbation-independent: a) increase in the variability of kinematics; b) increase in the relative duration of motor primitives; c) decrease in the accuracy of motor primitives. However, lesioned wild-type mice did not substantially modify any of the aforementioned parameters, indicating a major absence of adaptation that even led, in three out of five animals, to abrupt stops on the treadmill right after a sudden acceleration of the belt. Thus, our results reveal a key principle in supraspinal somatosensory processing: proprioceptive information from the DCML pathway is absolutely crucial to guarantee robust control of locomotion in challenging settings.

### Limitations and future directions

Lesions of the DCML pathway were carried out at the ninth thoracic segment. Thus, we cannot exclude that some dorsal spinocerebellar tract fibres were damaged, since it has been shown in the cat that the dorsal spinocerebellar tract neurons disappear completely from the dorsal column only at the sixth thoracic segment (English, 1985). However, also the dorsal spinocerebellar tract carries proprioceptive information from the hindlimb, thus our main goal of hindering ascending proprioceptive pathways would not be affected by a partial lesion of it. Moreover, both the DCML pathway and the dorsal spinocerebellar tract are known to additionally carry information from exteroceptors. Specifically, the DCML carries information from Aβ low-threshold rapidly-adaptive mechanoreceptors (Niu et al., 2013), while the dorsal spinocerebellar tract carries a mixture of touch and pressure signals (Hantman and Jessell, 2010; Lundberg, 1964). However, this should not substantially modify our conclusions, since locomotion has been shown to be largely independent of cutaneous feedback (Rossignol et al., 2006).

Stopping behaviours such those found in wild-type animals after lesion of the DCML pathway are still poorly understood, but there is strong evidence that the arrest of locomotor activity originates from supraspinal circuits (Roseberry and Kreitzer, 2017). In mice, the optogenetic activation of glutamatergic excitatory (Bouvier et al., 2015) or glycinergic inhibitory (Capelli et al., 2017) neurons in the caudal part of the brainstem has been proved to effectively suppress locomotion. Those caudal regions of the brainstem have synaptic interactions with the mesencephalic locomotor region that, if electrically or optogenetically stimulated, has been recently shown to arrest locomotion in the lamprey (Grätsch et al., 2019) and in the mouse (van der Zouwen et al., 2021), respectively. The supraspinal termination of movement after perturbation clearly illustrates the importance of stopping mechanisms as a defence strategy. Three conditions can lead to halting behaviour: goal completion, fear and startle (Roseberry and Kreitzer, 2017). With the current results, we cannot directly prove the origin of this behaviour and can only speculate that the animals in our experiment likely stopped due to fear and/or startle, rather than to complete a goal. This is nonetheless a further indication that the consequences of missing sensory afferent information were mediated by higher centres, resulting in the drastic interruption of the locomotor task.

### Conclusion

In conclusion, we showed that the systemic removal of proprioceptors in mice resulted in impaired locomotion proportionally to the level of ablation (i.e., of only muscle spindles or of both spindles and GTOs). This proves that sensory information from proprioceptors is needed to robustly cope with external perturbations and that the lack of sensory feedback has direct implications on the animal’s behaviour, negatively affecting movement and making muscle activation patterns less accurate (e.g., less complex) and adaptable. Moreover, the surgical interruption of ascending pathways carrying proprioceptive information to the brain through the spinal cord compromised the ability of wild-type mice to deal with perturbations, often leading to a stopping behaviour: a demonstration that supraspinal integration of sensory information from proprioceptors is crucial to robustly control locomotion at need.

## Materials and Methods

### Mouse lines

All procedures were performed according to the guidelines of the Canadian Council on Animal Care and approved by the local councils on animal care of Dalhousie University with protocols #19-014 and #19-021. Recordings were conducted on 17 adult (age 89 ± 26 days; mass 24.4 ± 2.9 g) male mice on C57BL/6 background fed ad libitum. Seven C57BL/6 wild-type mice were used for the DCML lesion protocol. To investigate the role of muscle spindles, we performed experiments with five *Egr3*^-/-^ mice (Tourtellotte and Milbrandt, 1998). To investigate the role of proprioceptive feedback from muscle spindles and Golgi Tendon Organs, we have used three *PV^Cre^::Avil^iDTR^* and two *PV^Cre^::Avil^iDTR^::Rosa^EGFP^* mice (Takeoka and Arber, 2019). Three *PV^Cre^::Rosa^EGFP^* mice were used for the sham experiments (Fig. S11). For the recordings, mice of different litters but same genotype were allocated to the relevant experimental group using as the only inclusion criteria: a) the genotyping, b) a minimum mass of 22 g and c) a minimum age of 50 days.

The mice used to breed the experimental mice were obtained from: *PV^Cre^*: Jackson Laboratories (017320) (Hippenmeyer et al., 2005); *Avil^iDTR^*: European Mouse Mutant Archives (EM: 10409) (Stantcheva et al., 2016); *Rosa^EGFP^*: Jackson Laboratories, USA (010701) (Sousa et al., 2009); *Egr3*^-/-^: courtesy of Dr. Warren Tourtellotte (Northwestern University, Chicago, IL; currently Cendar Cinai Medical Center, Los Angeles, CA); Wild type: littermates of different breeding colonies in the lab with C57/BL6 background.

### EMG implantation surgeries

Each mouse received an electrode implantation surgery as previously described (Akay et al., 2014; Santuz et al., 2019; Santuz and Akay, 2020). Eight bipolar EMG electrodes were implanted (Akay et al., 2006; Pearson et al., 2005) to as many muscles of the right hind limb. The muscles that were implanted were: gluteus maximus (MA), vastus medialis (VM), vastus lateralis (VL), semitendinosus (ST), biceps femoris (BF), tibialis anterior (TA), gastrocnemius medialis (GM) and gastrocnemius lateralis (GL). Briefly, mice were anesthetized with isoflurane at 2-3% concentration through inhalation at 1 l/min and ophthalmic eye ointment was applied to the eyes. The skin was sterilized by using a three-part skin scrub by means of Hibitane (Chlorhexidine gluconate 4%), alcohol and povidone-iodine. Two small midline skin incisions on the back and one on the posterior part of the right hind leg were made to expose the target muscles and position the custom 3D-printed back-mounted connector cap. The electrodes were then drawn from the most anterior neck incision to the leg incisions subcutaneously and the EMG connector cap was sutured to the skin. Afterwards, each electrode was implanted in the muscles as described before (Pearson et al., 2005), the incisions were closed, anaesthetic was discontinued and buprenorphine (0.03 mg/kg) and ketoprofen (5.00 mg/kg) were injected subcutaneously. The mice were then placed in a heated cage for at least six days and returned to their regular mouse rack once recovered. Food mash and hydrogel were provided for the first three days after the surgery multiple times a day and subsequently once a day until the animal returned to the rack. Handling of the mice was mostly avoided until they were fully recovered and at least for one week. Additional injections of buprenorphine and ketoprofen were performed in 24- and 12-hour intervals, respectively, for at least 48 hours after surgery.

### Spinal lesion surgeries

The DCML pathway lesion surgeries that were conducted after the first EMG recordings followed the same anaesthesia and care protocols as the EMG implantation surgeries. After preparation and anaesthesia, one midline skin incision of around 10 mm was made to expose the back muscles that were then separated with a scalpel to reveal the 7^th^ or 8^th^ thoracic vertebral body (Harrison et al., 2013). A laminectomy was then performed, the dura mater opened and the spinal cord lesioned at the 9^th^ (three mice) or 10^th^ (two mice) thoracic spinal segment using tungsten steel micro scissors. Muscles were then stitched over the missing spinous process using absorbable suture, the skin closed and the animals placed in their recovery cage for at least three days.

### Diphtheria toxin (DTX) delivery

Intraperitoneal DTX (D0654 lyophilized, Sigma-Aldrich, St. Louis, MO, United States) injection was carried out on three *PV^Cre^::Avil^iDTR^* and two *PV^Cre^::Avil^iDTR^::Rosa^EGFP^* experimental mice and on *PV^Cre^:: Rosa^EGFP^* mice for the sham experiments. We injected each mouse with 100 μg/kg of toxin (Takeoka and Arber, 2019) diluted in ultrapure water (Millipore, TMS-006-C) and let the animals in a confined cage for three days, when they could return to their rack. Recordings were conducted on the seventh day after injection.

### Immunohistochemistry

To assess the success of the DCML pathway lesion at the 9th/10th spinal segment, we perfused the animals immediately after euthanasia. After thoracotomy, the mice were perfused with 20 ml of saline solution followed by 10 ml of 4% paraformaldehyde solution (PFA) through the left cardiac ventricle. The spinal cord was dissected and immersed in PFA for at least 24 h. The 4th lumbar and 1st thoracic spinal segments were then dissected and cryoprotected by immersion in 30% sucrose-PBS solution for at least 24 h or until sunk. After cryoprotection, the spinal cord sections were embedded in optimal cutting temperature (OCT) mounting medium, flash frozen on dry ice and stored at −80 °C. The tissue was then sectioned transversally at 30 μm by means of a cryostat (Leica CM3050 S, Leica Biosystems AG, Muttenz, Switzerland) and placed on microscope slides to dry for at least one hour. Half of the sections were then prepared for Toluidine Blue O (TBO) staining and the remaining half for Protein Kinase C Gamma (PKCG) staining. For TBO (T3260 Sigma-Aldrich, St. Louis, MO, United States) staining, slides were washed three times for five minutes in 1xPBS to remove OCT and then immersed in 1:1000 TBO-1xPBS solution for other five minutes. Slides were then washed once for five minutes in 1xPBS and mounted on coverslips using PermaFluor Aqueous mounting medium (Thermo Fisher Scientific, Waltham, MA, United States). For PKCG staining of the corticospinal tract, after the three initial washes in 1xPBS, the slides were incubated for 60 minutes in 5% blocking solution (5% BSA in 1xPBS + 0.3% Triton) at room temperature. Subsequently, the sections were incubated overnight at 4 °C in a primary antibody solution consisting of 1:1000 rabbit-anti-PKCG (Abcam ab71558 rabbit pAB to PKCG, Abcam, Cambridge, UK) in 1% blocking solution (1% BSA in 1xPBS + 0.3% Triton). The next day, the tissue received three 10-minute washes in 1xPBS and a secondary incubation of 90 minutes at room temperature in 1:500 Invitrogen A32794 donkey-anti-rabbit alexa fluor 555 plus (Invitrogen, Carlsbad, CA, United States) in 1% blocking solution. After other three 10-minute washes in 1xPBS, slides were mounted on coverslips.

To assess the success of proprioceptive afferent ablation in *PV^Cre^::Avil^iDTR^* mutants after DTX injection, we dissected the dorsal root ganglia and spinal cord at the 4th lumbar spinal segment after perfusion. Following the procedures reported above, we prepared the tissue for Hoechst 33342 staining (both dorsal root ganglia and spinal cord), RUNX3 staining (dorsal root ganglia) and VGLUT1 staining (spinal cord). The procedure for Hoechst 33342 (Abcam ab228551) staining was identical to that described above for HBO staining. RUNX3 and VGLUT1 staining protocols were identical to that for PKCG, with different primary and secondary antibody incubations. For RUNX3 (Jessell laboratory), we incubated overnight at 4 °C with 1:5000 guinea pig-anti RUNX3 in 1% blocking solution and then for 45 minutes at room temperature with 1:1000 goat anti-guinea pig alexa fluor 555 plus (Invitrogen). For VGLUT1 (Jessell laboratory), we incubated overnight at 4 °C with 1:8000 rabbit-anti VGLUT1 in 1% blocking solution and then for 60 minutes at room temperature with 1:500 goat anti-rabbit alexa fluor 555 plus (Invitrogen).

### EMG recordings

After recovery from surgery, all animals walked on a custom-built treadmill (workshop of the Zoological Institute, University of Cologne, Germany) capable to administer perturbations through quick mediolateral displacements and accelerations of the belt. We recorded at least 30, often non-consecutive, gait cycles from each animal during unperturbed and perturbed walking. Each animal walked at two speeds: 0.25 and 0.30 m/s. In the case of severely compromised locomotor patterns, such as in the *PV^Cre^::Avil^iDTR^* mutants after DTX injection, the speed was reduced to 0.15 m/s to let the animals cope with the belt’s speed. Perturbations were administered at random time intervals (minimum 0 ms, maximum 1200 ms after the previous perturbation was completed) and consisted of either a) a sudden acceleration of the belt that doubled the speed and then returned to the initial one in 200 ms (100 ms acceleration, 100 ms deceleration) or b) a mediolateral motion that suddenly displaced the whole treadmill by 6 mm in 100 ms with constant velocity. The randomized perturbation protocol included 20 anteroposterior and 20 mediolateral perturbations, randomly distributed along 30 s (Fig. S1); the cycle started at random points and would repeat itself automatically if needed. A less challenging protocol, including only one mediolateral perturbation every 2 s, was administered to those animals that could not cope with the random protocol. Namely, this happened for the WT after DCML lesion. This reduced protocol produced similar effects to the randomized protocol on the gait spatiotemporal parameters (Fig. S2) and is referred to in the figures as “ramp”. EMG signals were recorded by connecting the electrodes via the EMG connector to an amplifier (MA 102, workshop of the Zoological Institute, University of Cologne, Germany) and recorded at a sampling frequency of 10 kHz. Animals were euthanized immediately after the last recording with a high dose of sodium pentobarbital (100 mg/kg) through intraperitoneal injection.

### Kinematics

The gait cycle segmentation (paw touchdown and lift-off timing) was obtained by the elaboration of high-speed videos. The kinematics of the hindlimb were acquired with a high-speed camera operating at 500 Hz with resolution 1280 × 800 pixels (IL3, Fastec Imaging, San Diego, CA, United States). For markerless body part tracking we used DeepLabCut (Mathis et al., 2018) v2.1.10. We labelled 16 landmarks on 297 frames taken from 30 videos of 15 different animals assigning the 95% of those images to the training set without cropping. Namely, we labelled six calibration markers, iliac crest and hip (highlighted by two white dots placed with an oil-based marker under brief 3% isoflurane anaesthesia through inhalation at 1 l/min), knee, ankle, fifth metatarsal, toe tip of the rear paw, toe tip of the front paw and the four paw reflections on the mirror placed under the treadmill at 45° angle. We used a ResNet-50-based neural network (He et al., 2016; Insafutdinov et al., 2016) with default parameters for 1030000 training iterations and two refinements of 100000 and 300000 iterations, respectively. We validated with one shuffle and found the test error was 2.75 pixels and the train error 2.57 pixels.

Of the 16 landmarks, we used eight for the segmentation of the gait cycle: the six calibration, the metatarsal and the rear toe tip markers. Following a procedure extensively reported previously (Santuz et al., 2018; Santuz and Akay, 2020), we processed the data to detect touchdown and lift-off of the right-side hindlimb. For touchdown estimation, we used the modified foot contact algorithm developed by Maiwald and colleagues (Maiwald et al., 2009). For estimating lift-off, we used the paw acceleration and jerk algorithm (Santuz et al., 2018; Santuz and Akay, 2020). We found [LOe – 20 ms, LOe + 20 ms] to be the sufficiently narrow interval needed to make the initial lift-off estimation. Both approaches have been validated against a set of 104 manually-labelled touchdown/lift-off events from 4 videos of two animals at different speeds (0.2 and 0.3 m/s) and showed a true error within the frame rate of the camera (i.e., ≤ 2 ms).

Poincaré maps were produced by mapping the hip-joint angle *θ_hip,norm_*, normalized between the minimum and the maximum of each gait cycle, at touchdown of the gait cycle n (x-axis) and at touchdown of the following gait cycle (y-axis). To quantify the dispersion of the obtained map, we used the descriptors (Brennan et al., 2001) SD1 and SD2, which are calculated as follows:

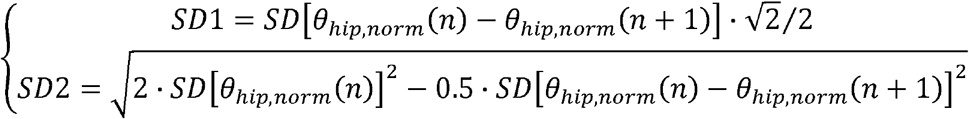

### EMG bootstrap

Given the challenges imposed by the implantation of many muscles (eight), some EMG channels suffered degradation of the signal between different sessions or right after surgery. In order to avoid issues in the extraction of muscle synergies originating from missing data, we used a bootstrap-like resampling approach reported before (Santuz et al., 2019). This allowed us to substantially reduce the number of needed animals, in accordance with the principles of humane animal research. Briefly, for each group and condition, we created 1000 resampled data sets each containing eight-muscle EMG data for 30 gait cycles. Each muscle activity in every gait cycle was randomly selected, without replacement, from all the available recorded activities for that group of animals and condition. In the case of perturbed locomotion, we followed the random perturbation protocol and inserted a perturbation every five steps or less (choosing randomly) and picking each time from the relevant data set (i.e., if the selected perturbation was a mediolateral displacement to the left, we would only pick from the EMG data sets that contained a mediolateral perturbation to the left for that cycle, etc.). We so obtained 1000 trials of 30 gait cycles each for WT, unperturbed walking, pre-DCML pathway lesion; another 1000 for *Egr3*^-/-^, perturbed walking; and so on for a total of 12000 trials, given the 12 analysed conditions.

### Muscle synergies extraction

Muscle synergies data were extracted from the bootstrapped EMG activity through a custom script (R v4.0.4, R Core Team, 2021, R Foundation for Statistical Computing, Vienna, Austria) based on the R package “musclesyneRgies” (Santuz, 2021) using the classical Gaussian nonnegative matrix factorization (NMF) algorithm as extensively reported in the past (Lee and Seung, 1999; Santuz et al., 2017). A schematic representation of the procedure can be found in Fig. 1. The R package “musclesyneRgies” for the pre-processing of raw EMG data and for the extraction and classification of muscle synergies is available at https://github.com/alesantuz/musclesyneRgies (Santuz, 2021). The source code version 0.7.1-alpha, used in this paper, is archived at Zenodo (Santuz et al., 2021).

### Fractal analysis of motor primitives

To assess the local and global complexity of the bootstrapped motor primitives, we calculated the Higuchi’s fractal dimension (HFD) and Hurst exponent (H), respectively (Higuchi, 1988; Hurst, 1951). The numerical procedures for obtaining both metrics were recently reported in detail (Santuz and Akay, 2020). HFD was calculated with a maximum window size of 10 points. H was calculated following the rescaled range (R/S) approach (Mandelbrot and Wallis, 1969) with a minimum window length of 200 points. HFD values range from 1 to 2, with increasing values correlating to increasingly complex data and HFD = 1.5 indicating random Gaussian noise (Anmuth et al., 1994; Higuchi, 1988; Kesić and Spasić, 2016). H can vary between 0 and 1. For 0.5 < H < 1, in the long-term high values in the time series (the motor primitive in our case) will be probably followed by other high values and a positive or negative trend is visible (Gneiting and Schlather, 2004; Mandelbrot, 1983). For 0 < H < 0.5, in the long term high values in the series will be probably followed by low values, with a frequent switch between high and low values (Gneiting and Schlather, 2004; Mandelbrot, 1983). H = 0.5 corresponds to a completely random series (Mandelbrot, 1983; Qian and Rasheed, 2004). In other words, values of H approaching 0.5 from both ends indicate more complex (or random) behaviour of the time series (Hurst, 1951).

### Sham experiments

Three *PV^Cre^::Rosa^EGFP^* mice underwent electrode implantation as described above. After recovery, the same recording protocol used for the other mice was performed. Then, the three animals were injected with DTX (Fig. S11) and after seven days another recording session was completed. Finally, the mice underwent the same surgery as the five lesioned animals, only this time without the actual lesion. The last recording session was then performed after recovery from surgery and the animals were perfused and dissected.

### Statistical analysis

To investigate the effects of perturbations and sensory ablation on the kinematics, gait spatiotemporal parameters, EMG activity and muscle synergy-related metrics, we followed a Bayesian multilevel modelling approach implemented in the R package *brms* 2.15.0 (Bürkner, 2017; Carpenter et al., 2017). For each variable of interest, we built a mixed effects model containing both “fixed” and “random” effects. The constant effects analysed were the locomotion condition (i.e., walking with or without perturbations), the animal group (with pre/post conditions where relevant) and their interaction. In all groups, we added the random effects as a by-animal varying intercept. Each model was run with five independent Markov chains of 10000 iterations, with the first 4000 warm-up iterations used only for calibration and then discarded, thus resulting in 30000 post-warm-up samples. Convergence of the chains and sufficient sampling of posterior distributions were confirmed by ensuring a potential scale reduction factor R□ < 1.01 and an effective sample size of at least 20% of the number of iterations. We so obtained the 95% credible intervals, defined as those intervals around the point estimates having a 95% probability of encompassing the population value, given the data and the prior assumptions (Nalborczyk et al., 2019). We used different priors depending on the investigated parameters and based on the mean and two standard deviations of previously recorded data. Specifically, we used normal priors with the following means and standard deviations: stance duration ~ N(115 ms, 55 ms); swing duration ~ N(95 ms, 30 ms); cadence ~ N(625 steps/min, 160 steps/min); Poincaré descriptors ~ N(0.2, 0.2); FWHM ~ N(65, 17); CoA ~ N(85, 30); H ~ N(0.55, 0.15). HFD values often produce target distributions having features with a resolution that cannot be grasped by the Hamiltonian model used by *brms*. Due to this, we used flat priors for HFD to avoid getting biased estimates. Since the EMG-related parameters (i.e., RMS_EMG_, FWHM, CoA, HFD and H) were obtained by resampling of the EMG signals as described above, the mixed effects models for these variables were obtained by sampling without replacement ten random values from the 1000 obtained with the bootstrap-like procedure. This was done in order to calculate the posterior distributions based on samples having the same size as the number of animals (five, two trials each) assigned to each group. Moreover, hindlimb joint angles were compared by evaluating the one-dimensional statistical parametric mapping (Pataky, 2012). All calculations were performed in R v4.0.4.

## Data availability

The datasets generated and analysed during the current study and an exhaustive explanation of their structure are available at the Zenodo repository (Santuz et al., 2021).

## Supplementary figures and movies

**Fig. S1.**
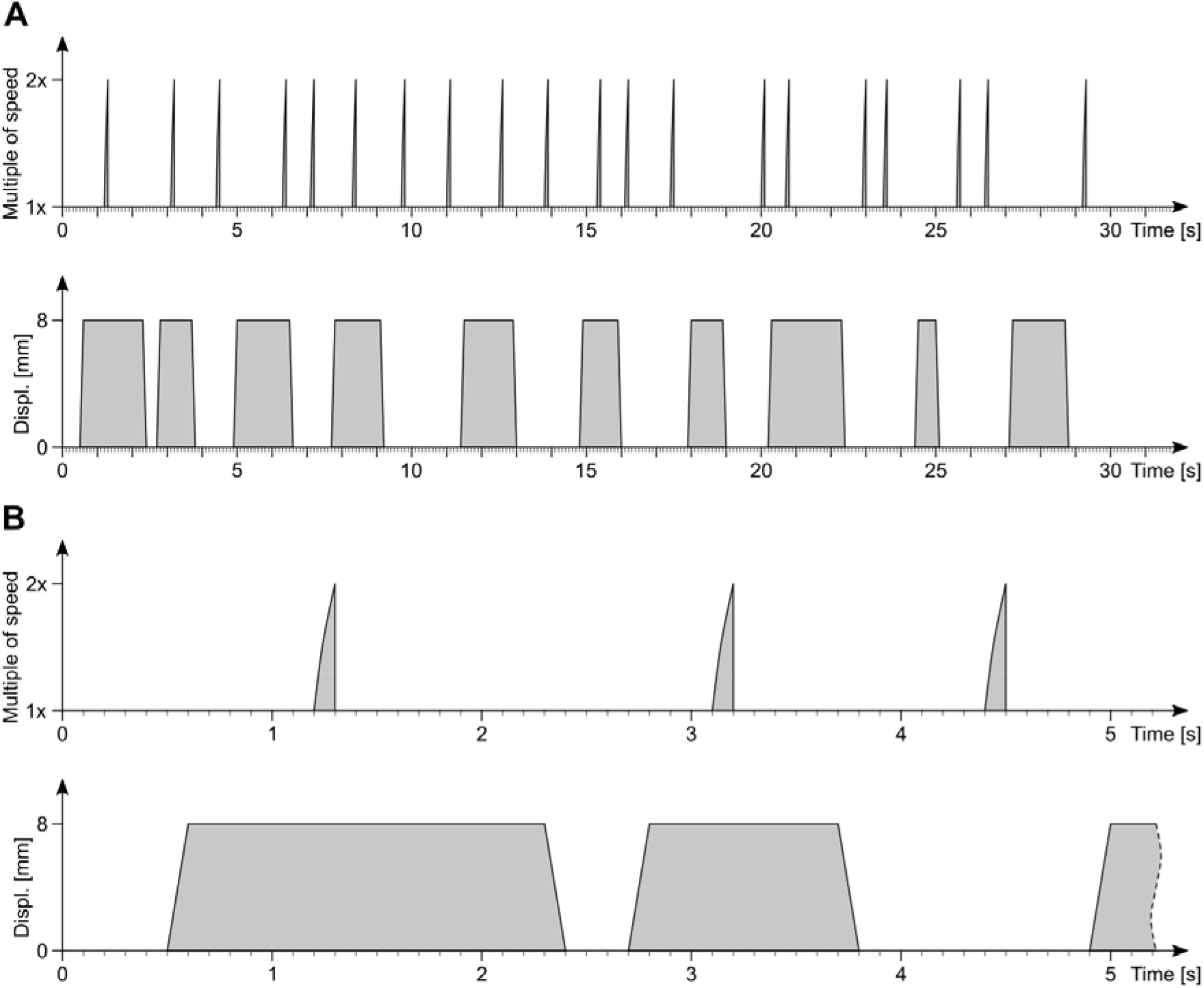
Schematic representation of the perturbation protocol. (**A**) The whole protocol of 30 s that would repeat itself indefinitely at need. Sudden accelerations (top) and mediolateral displacements (bottom) of the belt are presented as shaded areas. (**B**) Zoom in the first 5.2 s.

**Fig. S2.**
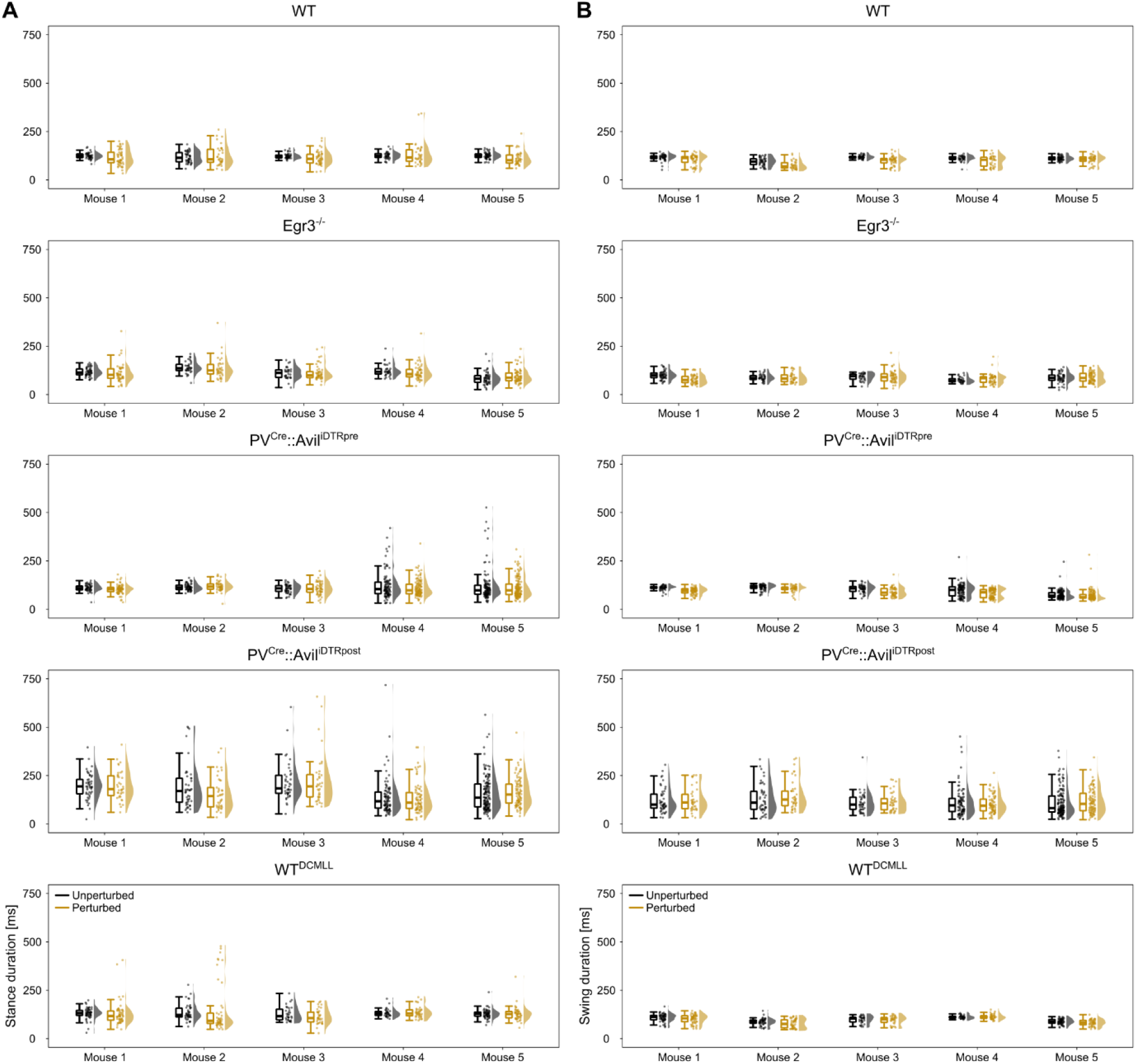
Gait temporal parameters. Stance (**A**) and swing (**B**) phase duration for each step recorded in every animal. Individual step values, pictured as dots and their distributions are shown next to the relevant boxplot.

**Fig. S3.**
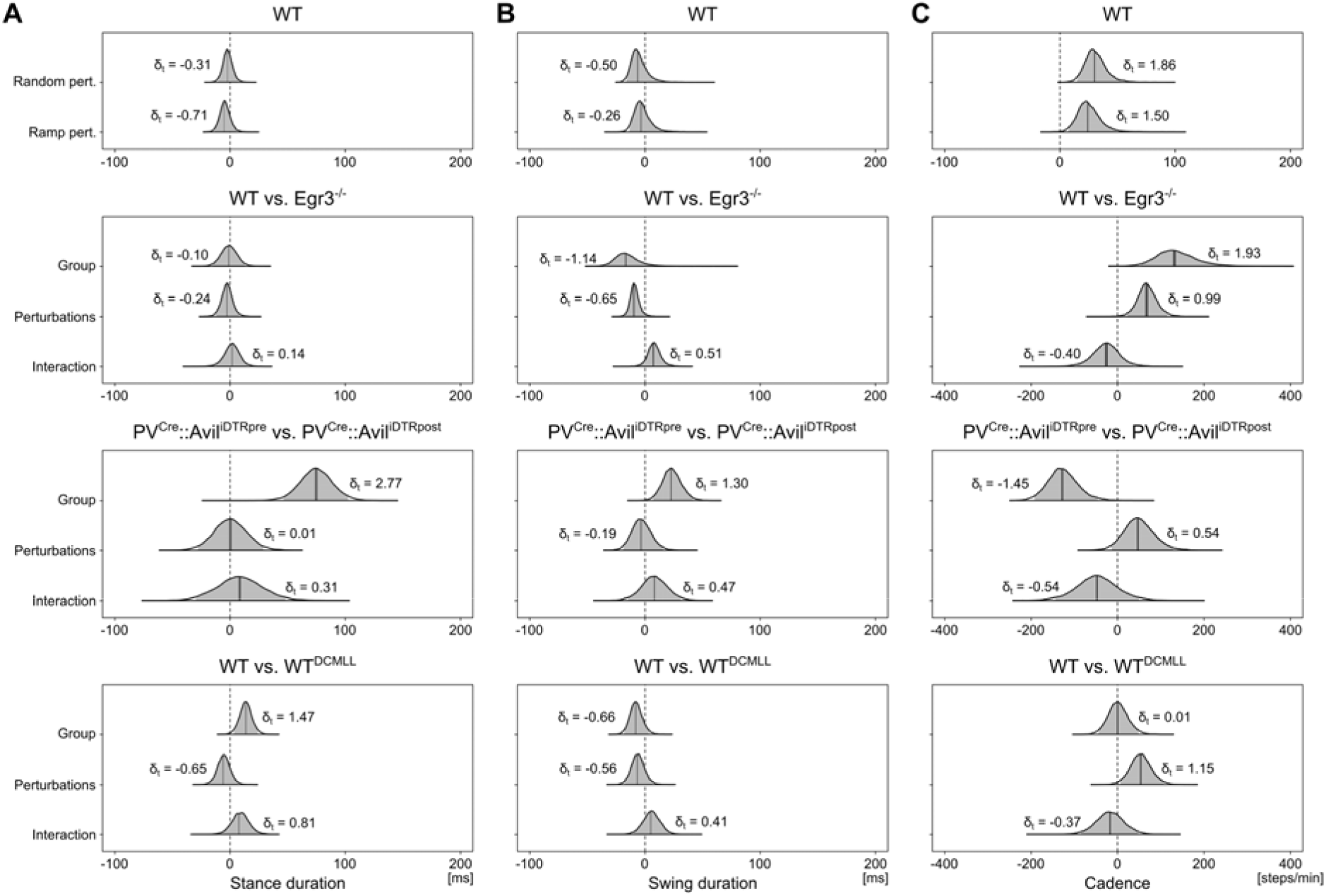
Posterior 95% credible intervals for the estimated differences in gait temporal parameters. (**A**) The 95% credible intervals and their probability distributions (shaded areas) describe the effects and interaction of the investigated groups and perturbations administered during locomotion on the stance duration. Effect size in the style of Hedges (i.e., considering all the variance sources in the model) are shown on the graphs and called δ_t_. The top-row graph shows a comparison between the standard perturbation protocol (“random”) and that including only one mediolateral displacement of the treadmill every 2 s (“ramp”). (**B**) The same as in panel (**A**), but for the swing phase duration. (**C**) The same as in panel (**A**), but for the cadence or number of steps per minute.

**Fig. S4.**
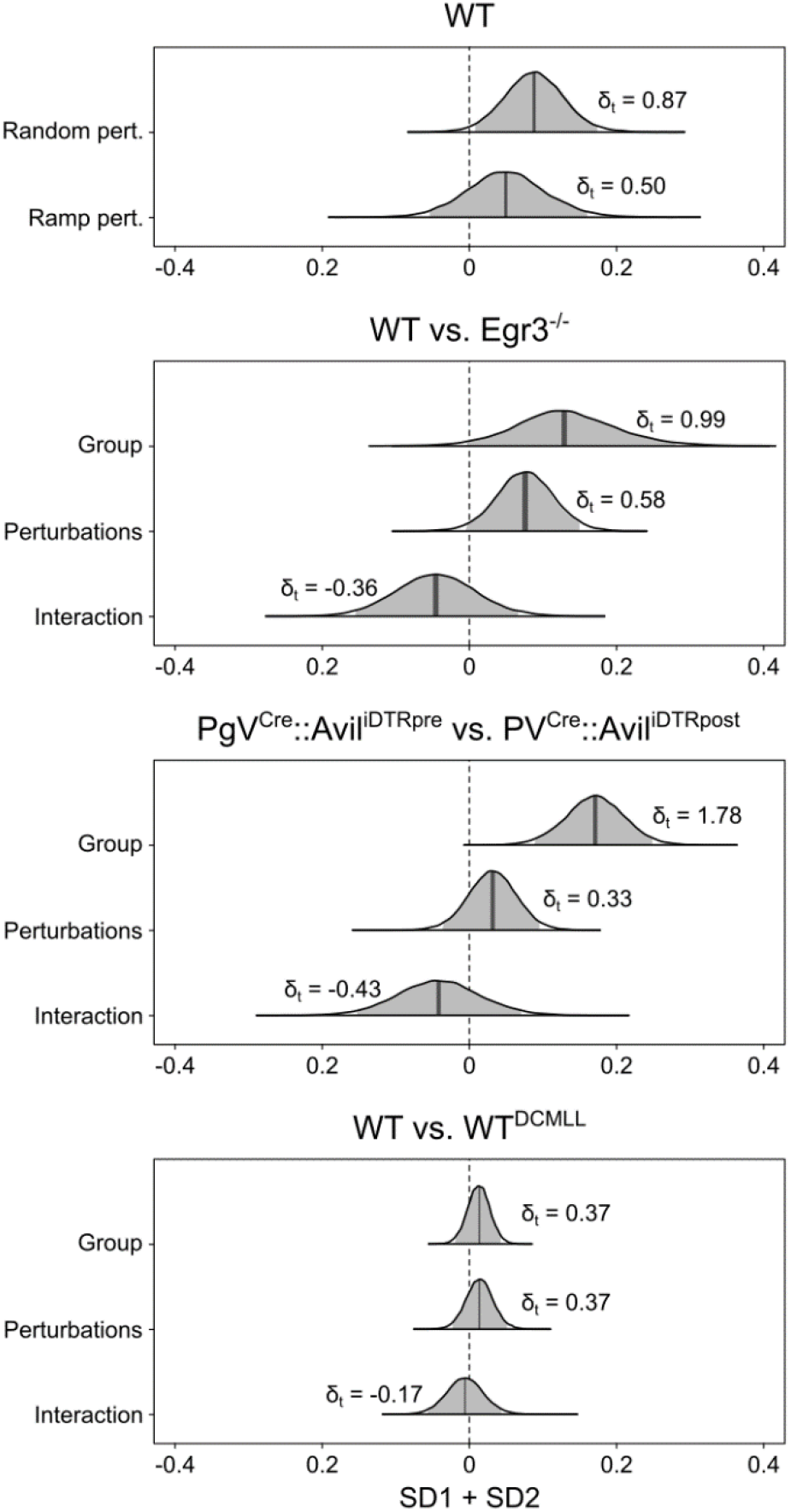
Posterior 95% credible intervals for the estimated differences in Poincaré map descriptors. The 95% credible intervals and their probability distributions (shaded areas) describe the effects and interaction of the investigated groups and perturbations administered during locomotion on the sum of the two Poincaré map descriptors (SD1 and SD2). The top-row graph shows a comparison between the standard perturbation protocol (“random”) and that including only one mediolateral displacement of the treadmill every 2 s (“ramp”). Effect size in the style of Hedges (i.e., considering all the variance sources in the model) are shown on the graphs and called δ_t_.

**Fig. S5.**
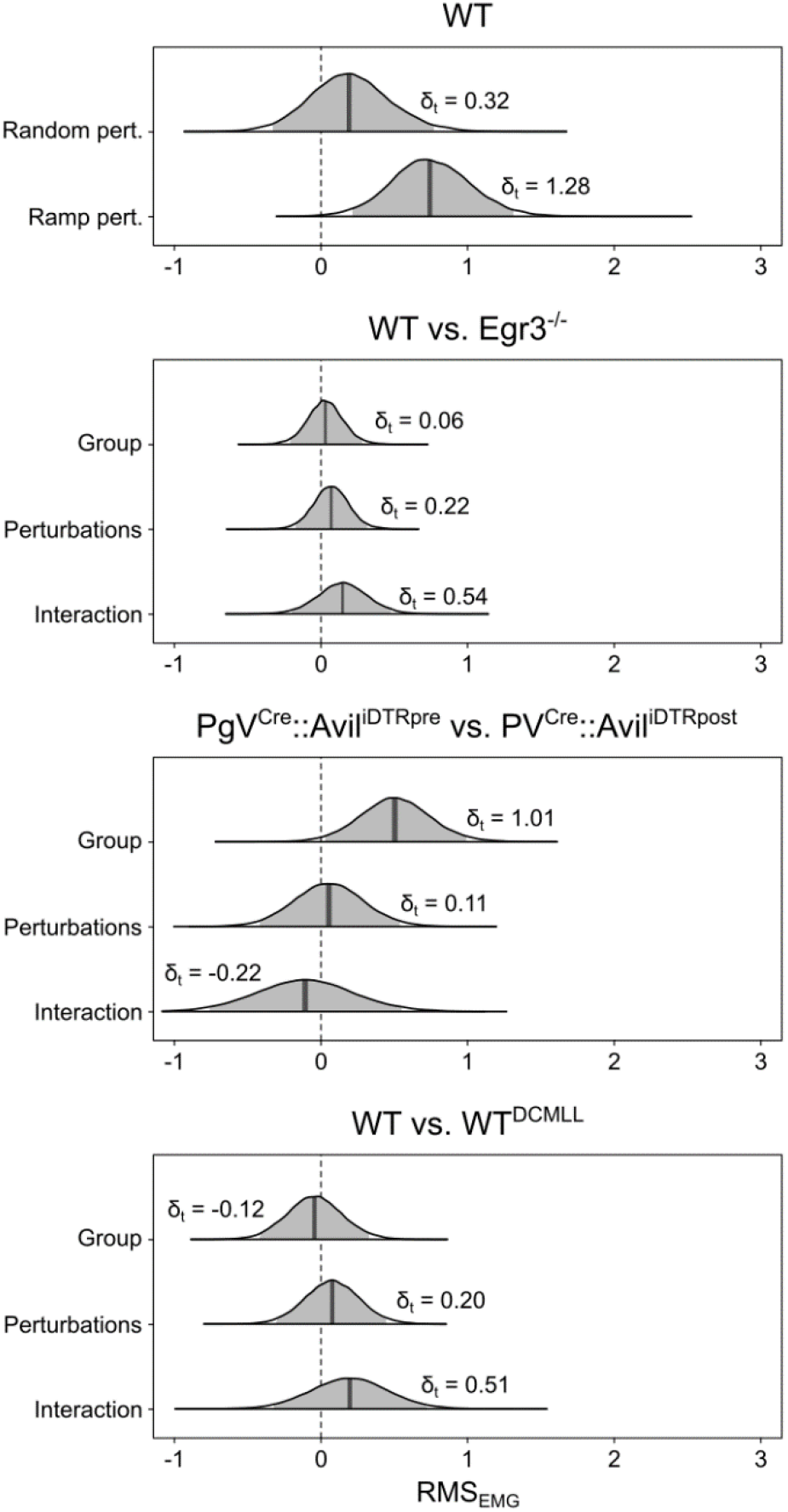
Posterior 95% credible intervals for the estimated differences in root mean square of the electromyographic signals (RMS_EMG_). The 95% credible intervals and their probability distributions (shaded areas) describe the effects and interaction of the investigated groups and perturbations administered during locomotion on the RMS_EMG_. The top-row graph shows a comparison between the standard perturbation protocol (“random”) and that including only one mediolateral displacement of the treadmill every 2 s (“ramp”). Effect size in the style of Hedges (i.e., considering all the variance sources in the model) are shown on the graphs and called δ_t_.

**Fig. S6.**
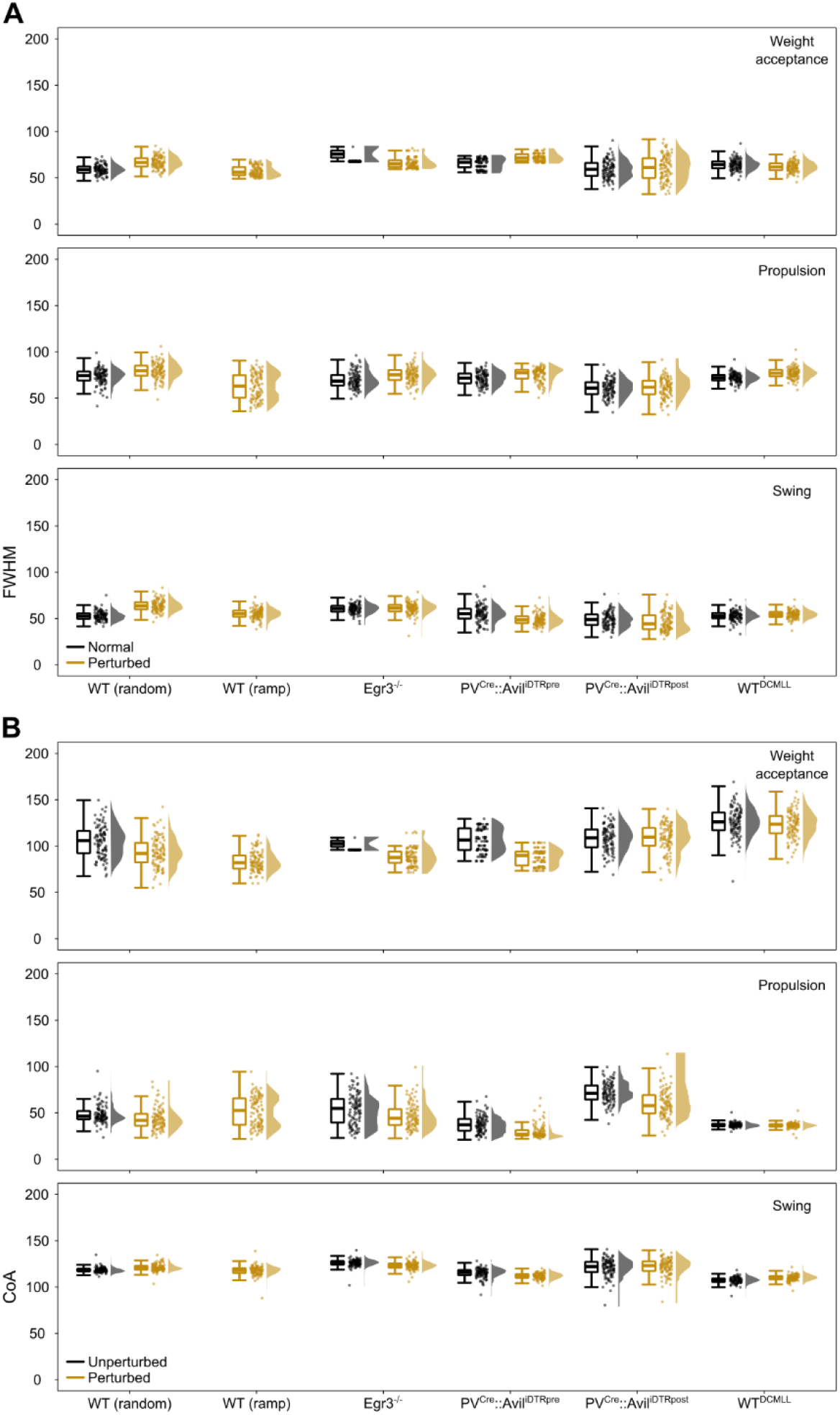
Full width at half maximum (FWHM) and centre of activity (CoA) of bootstrapped motor primitives. (**A**) Boxplots describing the FWHM of the bootstrapped motor primitives (weight acceptance, propulsion and swing) for the investigated groups. Raw data points (each point represents 10 nearest neighbours of the 1000 bootstrapped trials) and their density estimates are presented to the right side of each boxplot. (**B**) The same as in the previous panel, but for the CoA.

**Fig. S7.**
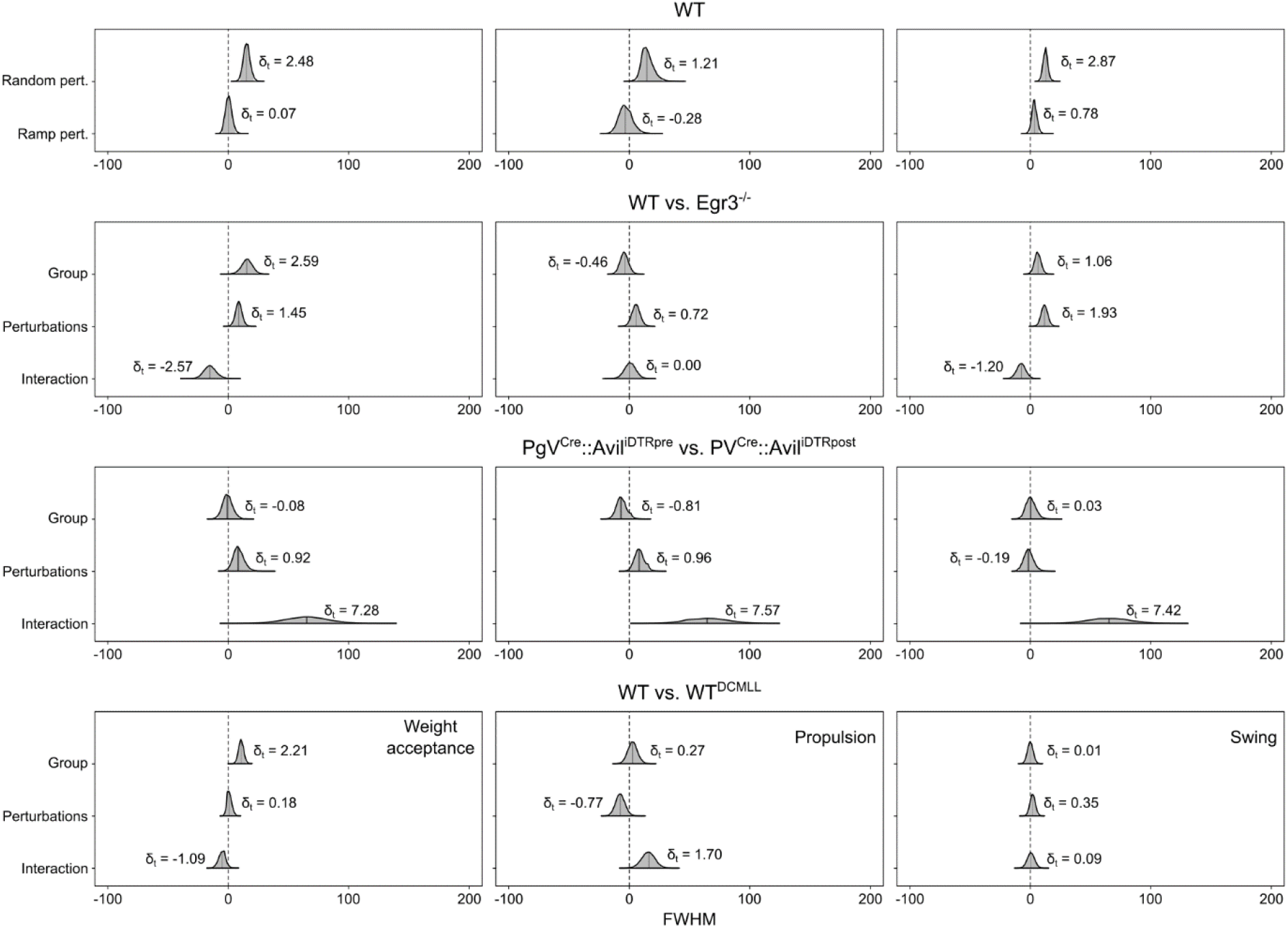
Posterior 95% credible intervals for the estimated differences in full width at half maximum (FWHM) of motor primitives. The 95% credible intervals and their probability distributions (shaded areas) describe the effects and interaction of the investigated groups and perturbations administered during locomotion on the FWHM of motor primitives (left: weight acceptance; middle: propulsion; right: swing). The top-row graphs show a comparison between the standard perturbation protocol (“random”) and that including only one mediolateral displacement of the treadmill every 2 s (“ramp”). Effect size in the style of Hedges (i.e., considering all the variance sources in the model) are shown on the graphs and called δ_t_.

**Fig. S8.**
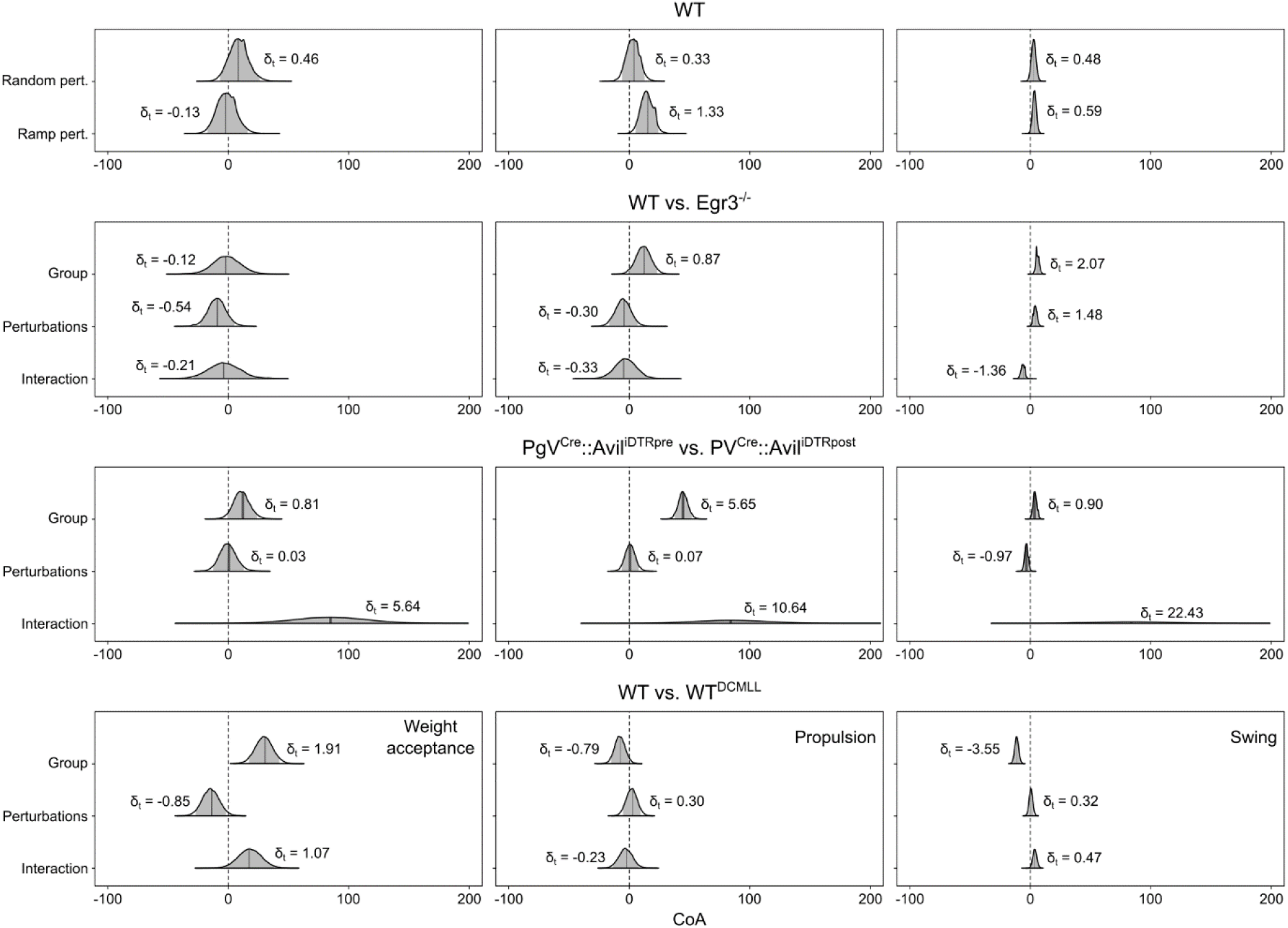
Posterior 95% credible intervals for the estimated differences in centre of activity (CoA) of motor primitives. The 95% credible intervals and their probability distributions (shaded areas) describe the effects and interaction of the investigated groups and perturbations administered during locomotion on the CoA of motor primitives (left: weight acceptance; middle: propulsion; right: swing). The top-row graphs show a comparison between the standard perturbation protocol (“random”) and that including only one mediolateral displacement of the treadmill every 2 s (“ramp”). Effect size in the style of Hedges (i.e., considering all the variance sources in the model) are shown on the graphs and called δ_t_.

**Fig. S9.**
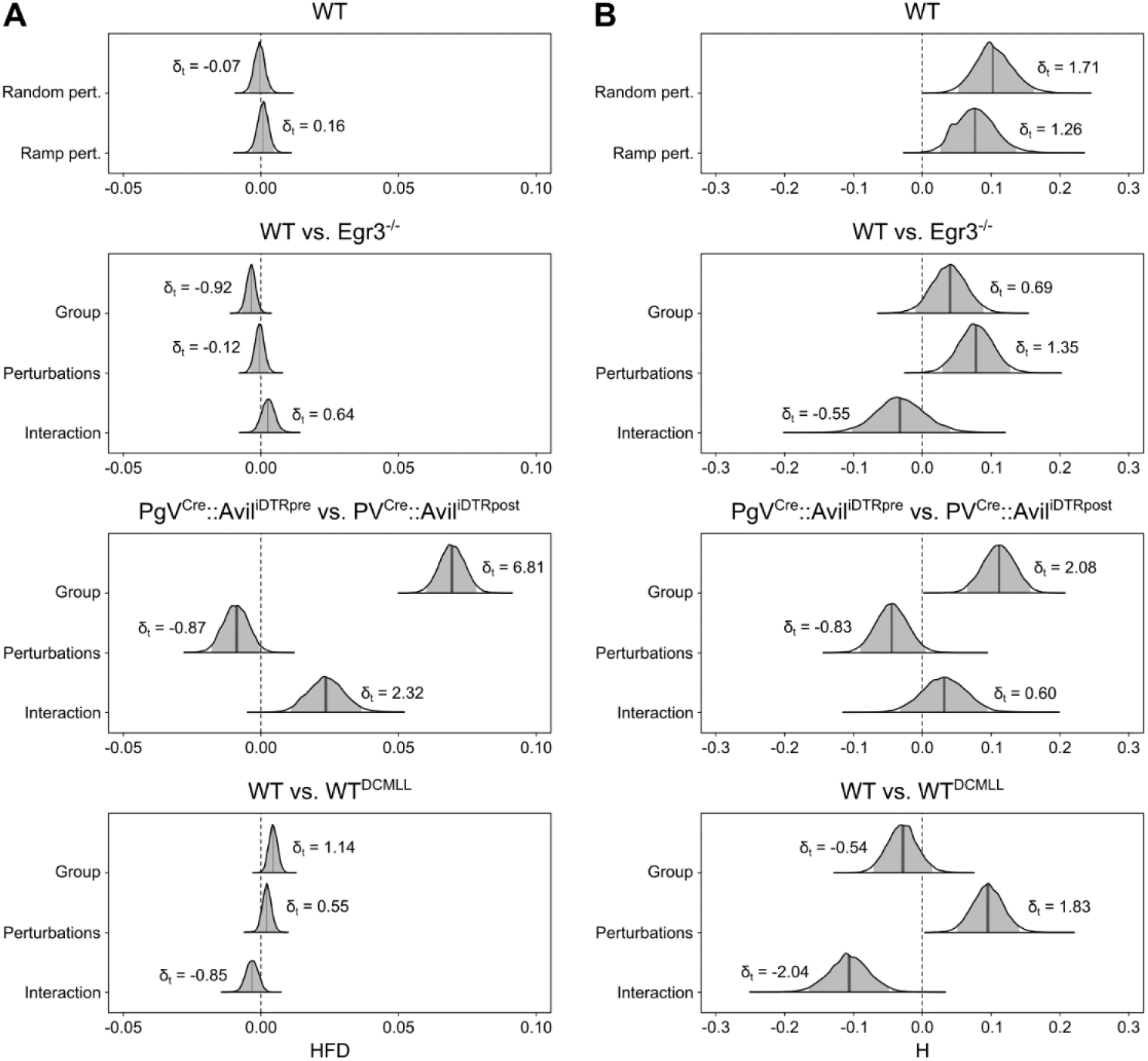
Posterior 95% credible intervals for the estimated differences in fractal dimension of motor primitives. (**A**) The 95% credible intervals and their probability distributions (shaded areas) describe the effects and interaction of the investigated groups and perturbations administered during locomotion on the Higuchi’s fractal dimension (HFD) of motor primitives. The top-row graph shows a comparison between the standard perturbation protocol (“random”) and that including only one mediolateral displacement of the treadmill every 2 s (“ramp”). Effect size in the style of Hedges (i.e., considering all the variance sources in the model) are shown on the graphs and called δ_t_. (**B**) The same as in the previous panel, but for the Hurst exponent (H).

**Fig. S10.**
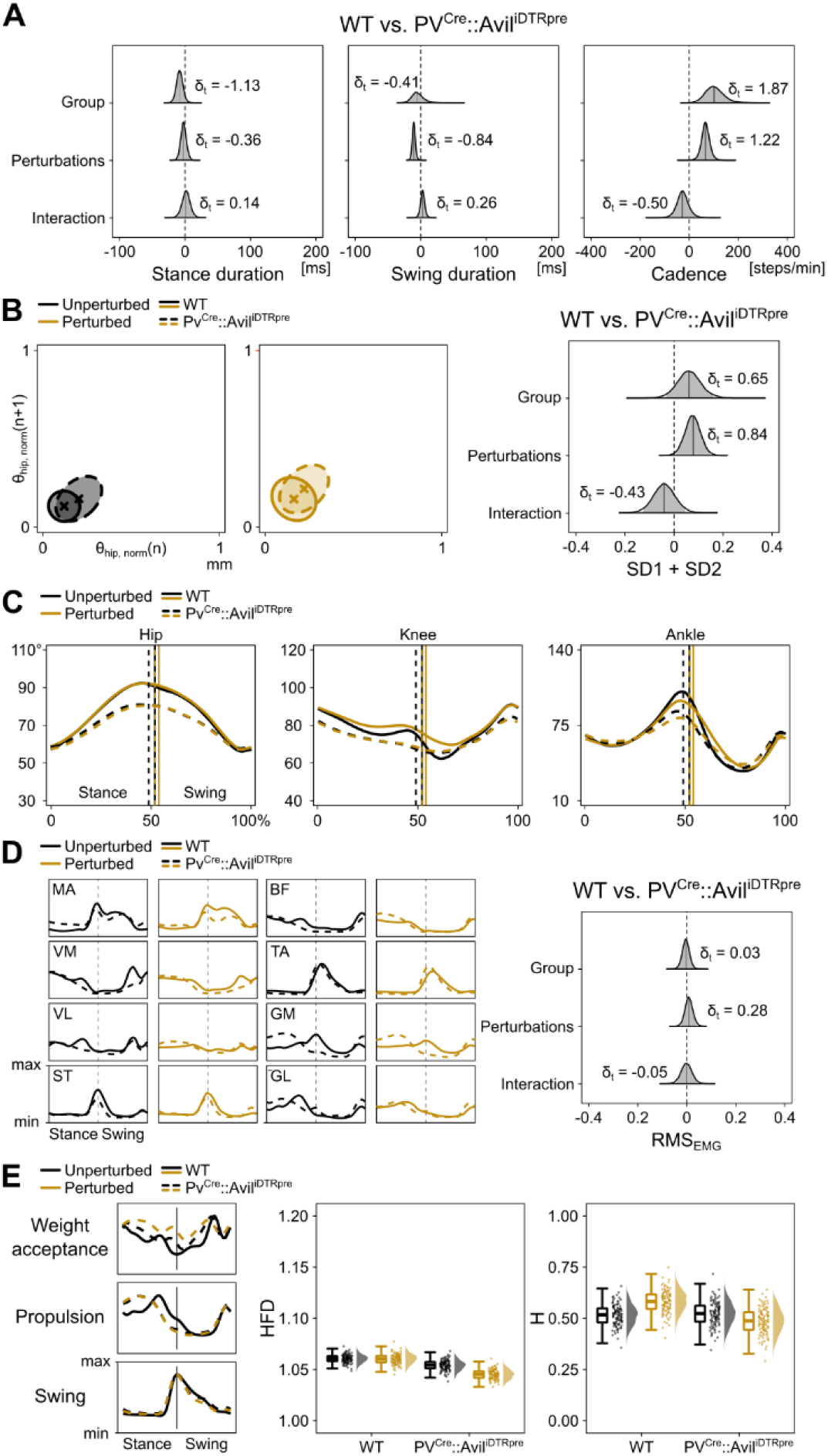
Largely similar behaviour of wild-type and *PV^Cre^::Avil^iDTR^* mice, before diphtheria toxin injection. (**A**) The 95% credible intervals and their probability distributions (shaded areas) describe the effects and interaction of the investigated groups and perturbations administered during locomotion on the stance duration (left), swing duration (middle) and cadence (right). Effect size in the style of Hedges (i.e., considering all the variance sources in the model) are shown on the graphs and called δ_t_. (**B**) Poincaré maps (left) of the hip angle θ_hip_ at touchdown (n) and (n+1) of all animals, with descriptor ellipse (see methods) and 95% credible intervals for the Poincaré descriptors. (**C**) Average joint angles. (**D**) Average electromyographic activity of the eight recorded muscles (left) and 95% credible intervals of the signal’s root mean square (RMS_EMG_). Muscle abbreviations: MA = *gluteus maximus*, VM = *vastus medialis*, VL = *v. lateralis*, ST = *semitendinosus*, BF = *biceps femoris*, TA = *tibialis anterior*, GM = *gastrocnemius medialis* and GL = *g. lateralis*. (**E**) Average motor primitives (left) and their complexity (right). Boxplots describe the Higuchi’s fractal dimension (HFD) and Hurst exponent (H) of the bootstrapped primitives. Raw data points (each point represents 10 nearest neighbours of the 1000 bootstrapped trials) and their density estimates are presented to the right side of each boxplot.

**Fig. S11.**
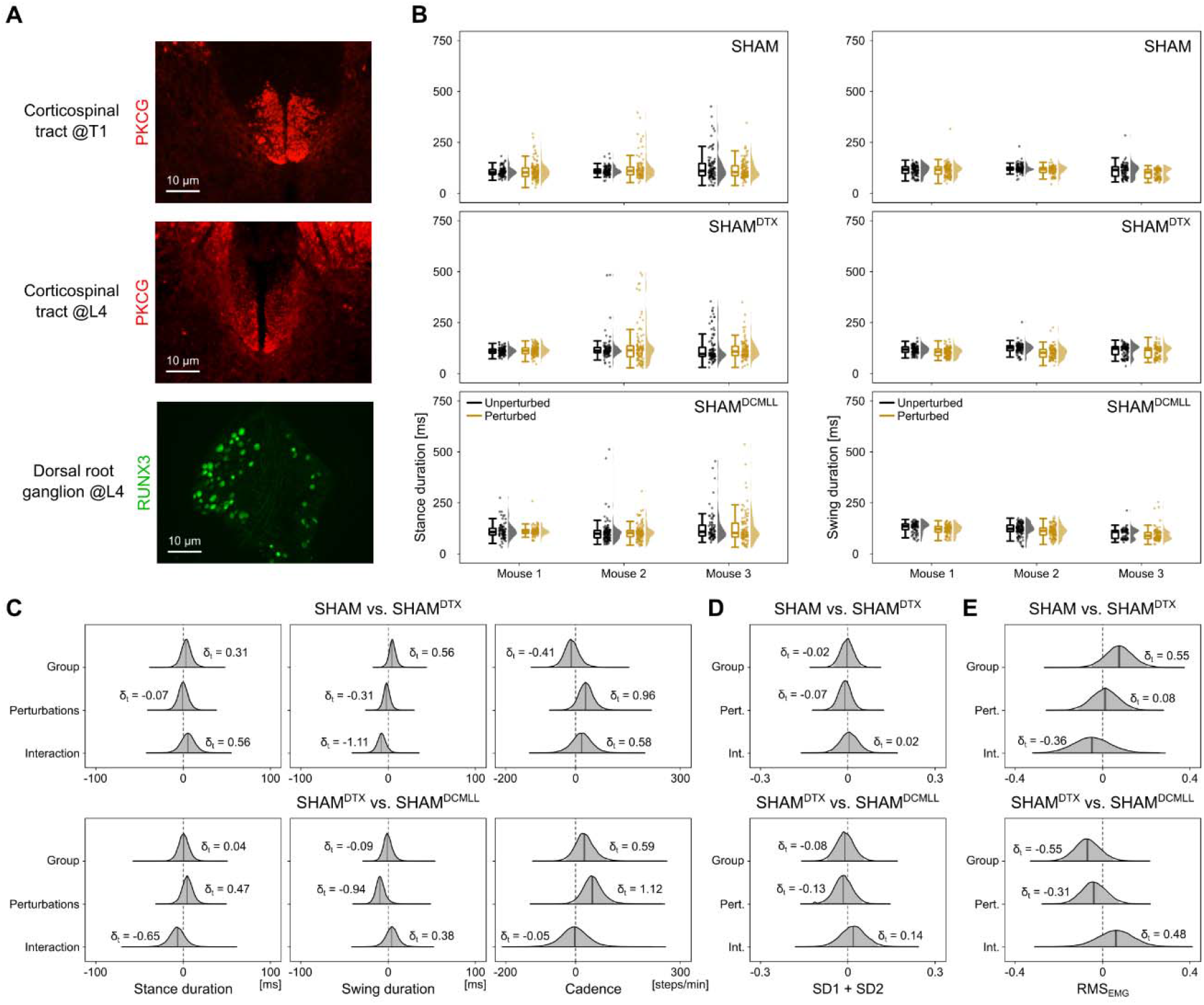
Sham experiments for diphtheria toxin (DTX) injections and dorsal column-medial lemniscus pathway lesion (DCMLL). (**A**) For the sham experiments, three animals underwent both DTX injection and DCMLL surgery (lesion excluded). Protein kinase C gamma (PKCG) is expressed in the axons of the corticospinal tract (CST). Here, both rostral (first thoracic spinal segment T1) and caudal (fourth lumbar spinal segment L4) sections with respect to the sham surgery site show intact CST. Runt-related transcription factor 3 (RUNX3) is selectively expressed in intact proprioceptive sensory neurons located in dorsal root ganglia. After DTX injection in sham mice that do not express human DTX receptors, the proprioceptive neuron terminals are left intact. (**B**). Stance (left) and swing (right) phase duration for each step recorded in every animal used for the sham experiments. Individual step values, pictured as dots and their distributions are shown next to the relevant boxplot. (**C**). The 95% credible intervals and their probability distributions (shaded areas) describe the effects and interaction of the investigated conditions and perturbations administered during locomotion on the stance duration (left), swing duration (middle) and cadence (right). Effect size in the style of Hedges (i.e., considering all the variance sources in the model) are shown on the graphs and called δ_t_. (**D**). The same as in (**C**), but for the two Poincaré map descriptors (SD1 and SD2). (**E**) The same as in (**C**), but for the root mean square of the electromyographic signals (RMS_EMG_).

**Movie S1.** Wild-type mouse walking on a treadmill while accelerations or mediolateral displacements of the belt are administered at random time intervals.

**Movie S2.** *Egr3*^-/-^ mouse lacking muscle spindles from birth walking on a treadmill.

**Movie S3.** *PV^Cre^::Avil^iDTR^* mouse after acute systemic ablation of muscle spindles and Golgi tendon organs walking on a treadmill.

**Movie S4.** Wild-type mouse after surgical lesion of the dorsal column-medial lemniscus pathway walking on a treadmill while accelerations or mediolateral displacements of the belt are administered at random time intervals.

## Acknowledgments

The authors are grateful to Brenda Ross for helping with the surgeries and maintaining the mouse colony, to Rachel Banks and Tyler Wells for supporting electrode production and tissue preparation, to Katrina Meyer for the immunohistochemistry suggestions and to Jacques Duysens and Sten Grillner for their comments on the manuscript. We also thank the Jessell lab, particularly Susan Brenner-Morton for sharing their primary antibodies against RUNX3 and VGLUT1 and Warren Tourtellotte for providing the Egr3-/- breeder mice.

## Author contributions

Conceptualization: A.S. and T.A.; Data curation: A.S.; Formal analysis: A.S.; Investigation: A.S. and O.D.L.; Software: A.S.; Supervision: T.A.; Visualization: A.S.; Writing – original draft: A.S.; Writing – review & editing: A.S., O.D.L. and T.A.

